# Contrasting selection at multiple life stages maintains divergent adaptation between sunflower ecotypes

**DOI:** 10.1101/2020.08.08.242503

**Authors:** April M. Goebl, Nolan C. Kane, Daniel F. Doak, Loren H. Rieseberg, Kate L. Ostevik

## Abstract

Conspecific populations living in adjacent, but contrasting, microenvironments represent excellent systems for studying natural selection. These systems are valuable because gene flow maintains genetic homogeneity except at loci experiencing strong, divergent selection. A history of reciprocal transplant and common garden studies in such systems, and a growing number of genomic studies, have contributed to understanding how selection operates in natural populations. While selection can vary across different fitness components and life stages, few studies have investigated how this ultimately affects allele frequencies and persistence of divergent populations. Here, we study two sunflower ecotypes in distinct, adjacent habitats by combining demographic models with genome-wide sequence data to estimate fitness components, absolute fitness, and allele frequency change at multiple life stages. This framework allows us to demonstrate that only local ecotypes experience positive population growth (lambda>1) and that the maintenance of divergent adaptation is mediated via habitat- and life stage-specific selection. We identify genetic variation, significantly driven by loci in chromosomal inversions, associated with different life history strategies in neighbouring ecotypes that optimize different fitness components and contribute to the persistence of each ecotype in its respective habitat.

## Introduction

Some of the most powerful studies documenting how selection operates in natural populations have focused on local adaptation of adjacent populations living in sharply divergent settings (Jain & Bradshaw, 1966; Schluter, 1993; Nosil et al. 2002; Sambatti & Rice, 2006). In these situations, gene flow has the potential to impede or reverse local adaptation, while habitat differences select for distinct sets of often co-varying traits. Multiple models have been developed to explain how strong selection can maintain divergence in the face of even high levels of genetic exchange (Maynard Smith, 1966; Caisse and Antonovics, 1978; Felsenstein, 1981). Most fundamentally, if selection coefficients are greater than the rate of migration, traits or regions of the genome under divergent selection will be maintained or continue to diverge, while unlinked features will not (Slatkin 1987; Wu 2001). Because only loci under divergent selection are expected to be differentiated, populations that are diverging with gene flow are valuable systems in which to study selection in nature.

Examples of such studies include Antonovics’ work on *Anthoxanthum* ecotypes on toxic mine tailings (McNeilly and Antonovics, 1968; Antonovics, 2006) and a long series of studies of serpentine and non-serpentine adapted ecotypes (Kruckeberg, 1950; Brady et al. 2005; Harrison & Rajakaruna, 2011). Traditionally, studies of these systems have used reciprocal transplant and common garden experiments to probe for phenotypic selection. This body of work has contributed to our understanding of how selection and gene flow interact to drive divergent adaptation in natural systems (Kawecki & Ebert, 2004; Sambatti & Rice, 2006; Ferris & Willis, 2018). More recently, researchers have used genomic methods to reveal patterns of selection by analyzing allele frequencies across habitat boundaries (Andrew & Rieseberg, 2013; Gompert et al. 2014; Egan et al. 2015; Tigano & Friesen, 2016). Using genomics has contributed to the study of divergent adaptation by addressing questions about the genomic architecture of ecologically important traits (Anderson et al. 2014; Hoban et al. 2016) and the temporal scale over which we expect selection to affect allele frequencies (Thurman and Barrett, 2016).

However, most phenotypic and genomic selection studies have not dissected the effects of selection on population fitness across different life history stages. Studies often focus on relationships between traits or genotypes and a single component of fitness, but this approach can potentially be misleading (Lande 1982; Cotto et al. 2019). Different fitness components can have different relationships to a given trait, and even the same fitness component can vary in its relationship to a given trait at different life history stages (Coulson et al. 2003; Ehrlen and Munzbergova 2009; Horvitz et al. 2010). In plants for example, seed size may be negatively correlated with survival at the seed stage (due to increased predation), positively correlated with survival at the seedling stage (due to higher resources), and negatively correlated with seed number at the adult stage. Additionally, each fitness component may be environmentally dependent, such that different life history strategies are favoured or selected against in different environments (Coulson et al. 2003; Cotto et al. 2019). Correctly integrating these fitness components across environments and life history stages in order to understand the effects on population persistence can be challenging.

Demographic analysis, the study of stage-specific growth, survival, and fecundity, can provide estimates of per capita population growth arising from multiple components of fitness. These estimates can be used to assess population dynamics in an ecological context, and as a measure of population fitness in an evolutionary context (Takada and Shefferson 2018). This approach to estimating trait effects on absolute fitness, as well as local adaptation, enables the question of habitat-specific persistence of individual ecotypes and hybrids to be addressed. The contribution of each fitness component to differences in ecotype fitness can also be determined. Therefore, simultaneously using genomic and demographic life history approaches presents a powerful framework. Yet studies of local adaptation seldom consider absolute fitness and we were unable to find any previous studies that combined demographic analyses modeling absolute fitness or population growth (sensu Caswell 2001) with genome-wide quantification of selection occurring in diverging populations.

Here we use genomic and demographic analyses to study rarely explored aspects of ecotypic divergence in an annual sunflower. In Great Sand Dunes National Park and Preserve in Colorado, USA neighbouring ecotypes of prairie sunflower (*Helianthus petiolaris*) inhabit two distinct habitats (Andrew et al. 2013). One unique ecotype has colonized the active sand dune system. Adaptation to the dunes is occurring despite opportunity for gene flow from the large, surrounding source ecotype that inhabits a vegetated sand sheet (Andrew et al. 2012). While the non-dune ecotype is typical for the species, the dune ecotype has some notable, genetically-determined phenotypic differences (Andrew et al. 2013; Ostevik et al. 2016). Previous research in this system showed indistinguishable allele composition across most of the genome between the two ecotypes, presumably due to the recent divergence and the ongoing homogenizing effects of gene flow, but also revealed several regions of significant genomic differentiation, suggestive of selection (Andrew and Rieseberg 2013). Recent studies identified that these regions of genomic differentiation correspond to large segregating inversions that are associated with several divergent phenotypes and habitat differences (Huang et al. 2020; Todesco et al. 2020). Additional work in this system has identified several partial barriers to gene flow between these ecotypes, both at pre- and post-zygotic stages (Ostevik et al. 2016).

While selection on multiple traits is important in these populations (Ostevik et al. 2016), it is not known how and at what life stages ongoing selection acts on individual adaptive alleles or distinct demographic traits. Previous studies illuminating genomic differentiation and loci associated with environment cannot discern when selection is acting on these loci. This is a powerful system to study contemporary allele frequency change because gene flow maintains variation within both populations, even for loci under selection. This means that selection can act on a wide variety of genetic combinations, in addition to the pure ecotypes. By analyzing genomic data from different life stages, we can assess when selection is strongest in each population.

We use this *H. petiolaris* system to link allele frequency shifts with measures of multiple components of fitness to investigate the maintenance of adaptive divergence between ecotypes. Using existing demographic data from a reciprocal transplant experiment (Ostevik et al. 2016) and new DNA sequence data from these plants, we ask i) how do fitness components that act at different life history stages contribute to overall fitness and persistence in each habitat? And ii) how does selection at different life history stages change allele frequencies, both genome wide and in putatively adaptive inversions, in each habitat? Studying local adaptation by separating selection on different traits and their environment-specific fitness effects on allele frequencies can inform how selection for ecotypic differences occurs across the life cycle, and how these effects may magnify or dampen overall divergence.

## Materials and Methods

### Study system

Great Sand Dunes National Park and Preserve (GSD) located in Southern Colorado, USA (37.7916°N, 105.5943°W) contains two ecotypes of *H. petiolaris*. One inhabits the active sand dunes (hereafter dune ecotype), and one grows adjacent to the dunes in prairie, non-dune habitat (hereafter non-dune ecotype); populations of dune and non-dune can be separated by as little as 100m (Andrew et al. 2012). *Helianthus petiolaris*, native to North America, is an outcrossing, hermaphroditic Compositae (Asteraceae) species that typically inhabits dry prairies and partially sandy soils. Dune individuals experience less stable soil (effectively 100% sand), lower soil nutrients, and less vegetation cover compared to the non-dune habitat (Andrew et al. 2012). While the non-dune ecotype is phenotypically and genetically typical for the species, the dune ecotype is characterized by large seeds (>2x larger than non-dune seeds), rapid seedling growth, reduced branching, and thicker stems (Andrew et al. 2013; Ostevik et al. 2016). In GSD the average monthly temperature during the growing season (April - October) is 13.3°C (based on GSD Remote Automatic Weather Station data averaged from 2005-2012), and the average cumulative growing season precipitation is 222.7mm (based on GSD National Weather Service station data averaged from 1950-2009). The average monthly temperature during the growing season and the cumulative growing season precipitation in the year of the field experiment (2012), were 14.4°C and 184.9mm, respectively.

### Reciprocal transplant experiment

Details of the 2012 reciprocal transplant experiment are published in Ostevik et al. (2016). Briefly, *H. petiolaris* seeds were collected from three dune and three non-dune populations at GSD in 2010. Seeds from these populations were used to generate F1 and backcross hybrids under greenhouse conditions: F1 offspring with both dune (F1D) and non-dune (F1N) cytoplasms, and backcrosses using both dune (BCD) and non-dune (BCN) pollen and equal proportions of dune and non-dune cytoplasms. A portion of the seeds from each population and each hybrid type (except BCN due to insufficient seed numbers) were grown in optimal greenhouse conditions and leaf tissue was collected for DNA sequencing (hereafter pre-selection samples). The remaining seeds from dune, non-dune and hybrid sources were planted in the field at GSD in cleared plots at one site in the dune habitat and one site in the non-dune habitat. Ten seeds from each population and hybrid type were planted per plot in a total of 45 plots per habitat. The following data were collected; seedling emergence (represents germination and early seedling survival), survival (represents survival from seedling to reproductive maturity), and number of seeds produced (represents total number of viable seeds per plant). Leaf tissue was collected for DNA sequencing from all plants that survived to flower (hereafter post-selection samples). Due to low emergence and survival during the experiment for some seed types, and resulting low sample sizes (table S1), we pooled plants based on source (dune, non-dune, or hybrid) for the following analyses. This means that our results for hybrids are based on a combined pool of individuals that included the different hybrid types (F1s and backcrosses). For our genetic analyses, we randomly sampled individuals of each hybrid type for the pre-selection pool to have proportions equal to the number of each type planted in the field (Supplemental Methods).

To account for the possibility of natural recruitment in experimental plots, we identified and removed any suspected local volunteers from the data set by examining the weight of seeds that plants produced (seed size is maintained in common gardens). Plants labelled as survivors in their foreign habitat that produced seeds outside of their ecotypic seed weight range and inside the range of the local ecotype (95% confidence intervals; Ostevik et al. 2016) were assumed to be local recruits and excluded from analyses (3 individuals; fig S1).

### Analysis of demographic data

All analyses of demographic data were done using R version 3.5.0 (R Core Team 2018) and followed general procedures in common use for demographic modeling (Caswell 2001, Morris and Doak 2002), applied to the separate stages of an annual life cycle (e.g. Smith et al. 2005, Griffith 2010).

#### Fitness component models

Data for seedling emergence, seedling-to-adult survival, and fecundity were modeled with source (i.e. dune, non-dune, or hybrid) and habitat of reciprocal transplant site (i.e. dune or non-dune) and their interaction as predictor variables. A binomial generalized linear model (GLM) with a logistic link function was fit to emergence data; this captures the probability that an individual seed germinated, emerged, and survived until the census (six weeks after planting). A binomial GLM was fit to survival data, which reflects the probability that a seedling survived to reproduce. Fecundity was determined by the number of seeds produced as follows; i) the number of viable-looking seeds in all collected heads were counted. ii) To adjust for heads that went missing prior to collection, average seed number per head was multiplied by the total number of inflorescences per plant, as determined by counts made throughout the flowering period. iii) Seed number was adjusted by removing the proportion of seeds observed to be eaten by insect larva. Fecundity therefore represents the total number of viable seeds produced that survived pre-dispersal predation. Number of seeds was then fit using a negative binomial model with a log link function.

Models including population of origin and planting plot as random effects were also fit to each fitness component. These models yielded similar results to those using only source and transplant habitat (table S2); we present results from the simpler models here.

#### Population model

To estimate lambda (per capita annual population growth rate, or fitness) for each source in each reciprocal transplant habitat, we multiplied fitness component estimates (emergence (E), survival (S), fecundity (F), seed survival (D)) that span the complete annual life cycle as follows: Lambda = ESFD. There was no data obtained for seed survival from this system, so we used a constant, biologically realistic rate (0.3 for all sources in both habitats) based on reports from the literature of other wild *Helianthus* populations (Alexander et al. 2001; Dechaine et al. 2010). Seed survival reflects the probability that a seed lands on suitable habitat, survives fungal attack, post-dispersal predation, and any other source of mortality occurring between dispersal in the fall and germination the following spring. We investigated how sensitive our estimates of lambda are to a range of seed survival values; we found that a broad range of values, ranging from 21% - 58%, yielded qualitatively similar results. To assess the error associated with our estimates of lambda we used a parametric bootstrapping approach: 10,000 simulations were obtained using parameter values randomly selected from multivariate normal distributions characterizing the parameter estimates for each fitness component. For each set of random parameter draws, we then calculated relative lambda (log ratio) between all source comparisons in both habitats and used 95% quantiles to determine significant differences between sources.

#### Life table response experiment

To quantify the contribution of each fitness component to differences in lambda between the two ecotypes growing in the same habitat, we used a life table response experiment (LTRE; Caswell 2001). The magnitude and direction of a given contribution in each habitat was assessed as the product of the sensitivity (partial derivative of a given fitness component to lambda while other fitness components were held at mean values) and the change in the given fitness component between the local and foreign ecotype (local – foreign). Error on estimates was determined using parametric bootstrapping and 95% quantiles (fig S3). LTREs were not conducted for hybrids. Differences in fitness between ecotypes in both habitats was quantified as the local ecotype lambda estimate minus the foreign ecotype lambda estimate.

### Genome sequence analysis

#### Genotyping-by-sequencing

Genomic DNA was extracted from leaf tissue from 437 individuals (250 pre-selection, 103 post-selection, and 84 from non-experimental adult plants (Supplementary Methods)) using a modified CTAB protocol (Doyle and Doyle 1987); reduced representation libraries were then prepared and sequenced (Supplementary Methods).

#### Sequence data processing

After quality control and trimming (Supplementary Methods), sequence reads were aligned to the *Helianthus annuus* reference genome Ha412HOv2 (Badouin et al. 2017) using bwa mem (Li 2013). Over 11.9 ×10^6^ variants were called and output in vcf format using samtools mpileup with minimum mapping quality cutoff (-q flag) of 20 and bcftools call with the multiallelic-caller model (Li 2011). Variants were filtered to only keep sites from single copy regions of the genome. Single copy sites were estimated by assessing the distribution of marker depth across the Ha412HO reference genome; >7.4 ×10^6^ variants remained after this filter. VCFtools (Danecek et al. 2011) was then used to filter variants for single nucleotide polymorphisms (SNPs) only (indels removed), biallelic sites, minimum mapping quality of 50, and a minor allele frequency cutoff of 5%; this resulted in 680,100 remaining SNPs. Genotypes were treated as missing if individual read depth was below 5 or genotype quality was below 20. Finally, SNPs were filtered iteratively based on individual and site missingness (table S3). Our final dataset contained genotype information for 411 plants at 12,214 SNPs distributed across the genome.

#### Analysis

The following analyses were done using R version 3.5.0 (R Core Team 2018). Principal component analysis (PCA) was performed using the R package PCAdapt (Luu et al. 2017) version 4.1.0 with K=5 and default arguments. Samples were grouped by source for this analysis. We used individual and mean PC scores to visualize the genetic structure of pre-selection samples (mean hybrid score was based on sampling individuals in proportion to the number of each hybrid type planted in the field; Supplementary Methods), and to test the effect of emergence and survival on post-selection allele frequencies in both habitats.

We investigated the genomic response to selection occurring at early life stages, and whether allele frequency change occurred at locations suspected to be targets of divergent selection. For this analysis we focused on the dune habitat where selection was strongest during early life stages. We first investigated hybrid individuals, for which we expect the most potential for evolutionary change and compared these results to the genomic response of dune samples. We excluded samples of non-dune source due to low post-selection numbers (table S1).

We calculated allele frequency change during early life stages by first calculating allele frequencies for each group (e.g. pre-selection hybrid source) and ecotype at every SNP as the sum of genotype calls across samples of a given group (n) divided by the number of chromosomes with data for a given SNP (n x 2). Next, allele frequency change of a given source was calculated by subtracting the frequency of the reference allele (arbitrarily selected over the alternate allele) in pre-selection samples from the frequency of the reference allele in post-selection samples. We also assessed allele frequency change by modeling allele count (reference vs alternate), for each source, as a function of selection (pre-vs post-) using a binomial GLM with a logistic link function at each SNP. We extracted p-values to assess the effect of selection on alleles. These two methods produced similar results (fig S10); we present data from the former method since units of allele frequency are more interpretable.

We investigated the relationship, via Pearson correlation, between allele frequency change in each source and allele frequency difference between ecotypes at each SNP. We quantified ecotype differences using the frequency of the reference allele in non-dune samples (non-experimental adult plants from two non-dune populations) subtracted from the frequency of the reference allele in dune samples (non-experimental adult plants from one dune population). Significance was determined using one-sided tests and distributions from randomization tests (details in fig S7 caption).

We also investigated how loci in seven genomic regions that likely contain inversions and locally adapted alleles responded during the experiment compared to sites across the rest of the genome. These putative inversions have been identified as highly differentiated between ecotypes and associated with traits and environmental features that differ between ecotypes (Huang et al. 2020; Todesco et al. 2020). We compared the distribution of allele frequency change in each inversion to the distribution at all non-inversion loci using a two-sample linear rank test with default arguments in the R package EnvStats v2.3.1 (Millard, 2013). Additionally, we tested allele frequency change in each inversion, when treating each inversion as a single locus (Huang et al. 2020; Supplementary Methods) using a Fisher’s exact test in the R package stats v3.6.2.

Lastly, to estimate the effects of fecundity on allele frequencies (in the absence of actual genotype data from the next generation), we weighted individual PC1 scores by the number of seeds produced by each plant. These weighted PC scores represent the estimated genotypes of seeds in the next generation. Samples from different sources were pooled for this analysis to increase variation in fecundity and genotypes for assessing any effects of differential fecundity on allele frequencies. Weighted PC scores were averaged across all samples in each pool: dune habitat (all dune, non-dune and hybrid plants that survived in the dune habitat), and non-dune habitat (all dune, non-dune and hybrid plants that survived in the non-dune habitat). In order to assess how allele frequencies changed in each habitat due to differential fecundity, weighted PC scores were compared to the mean PC score of pre-selection samples (dune, non-dune, and hybrid plants grown in greenhouse conditions and randomly sampled based on the proportion of each source planted in the field), and unweighted mean PC scores of dune habitat samples (all plants that survived in the dune habitat), non-dune habitat samples (all plants that survived in the non-dune habitat).

## Results

### Fitness components and population growth

We saw strong differences in the effects of habitat on plants from different sources at different life stages largely in line with Ostevik et al. (2016) (fig 1, table S4A-C). Dune and hybrid plants emerged and survived better in the dune habitat (although more dune plants emerged in the dune habitat than hybrid plants) but produced comparable numbers of seeds in both habitats. In contrast, non-dune plants emerged and survived at similarly low rates in both habitats but produced close to two orders of magnitude more seeds in their home environment.

**Figure 1:**
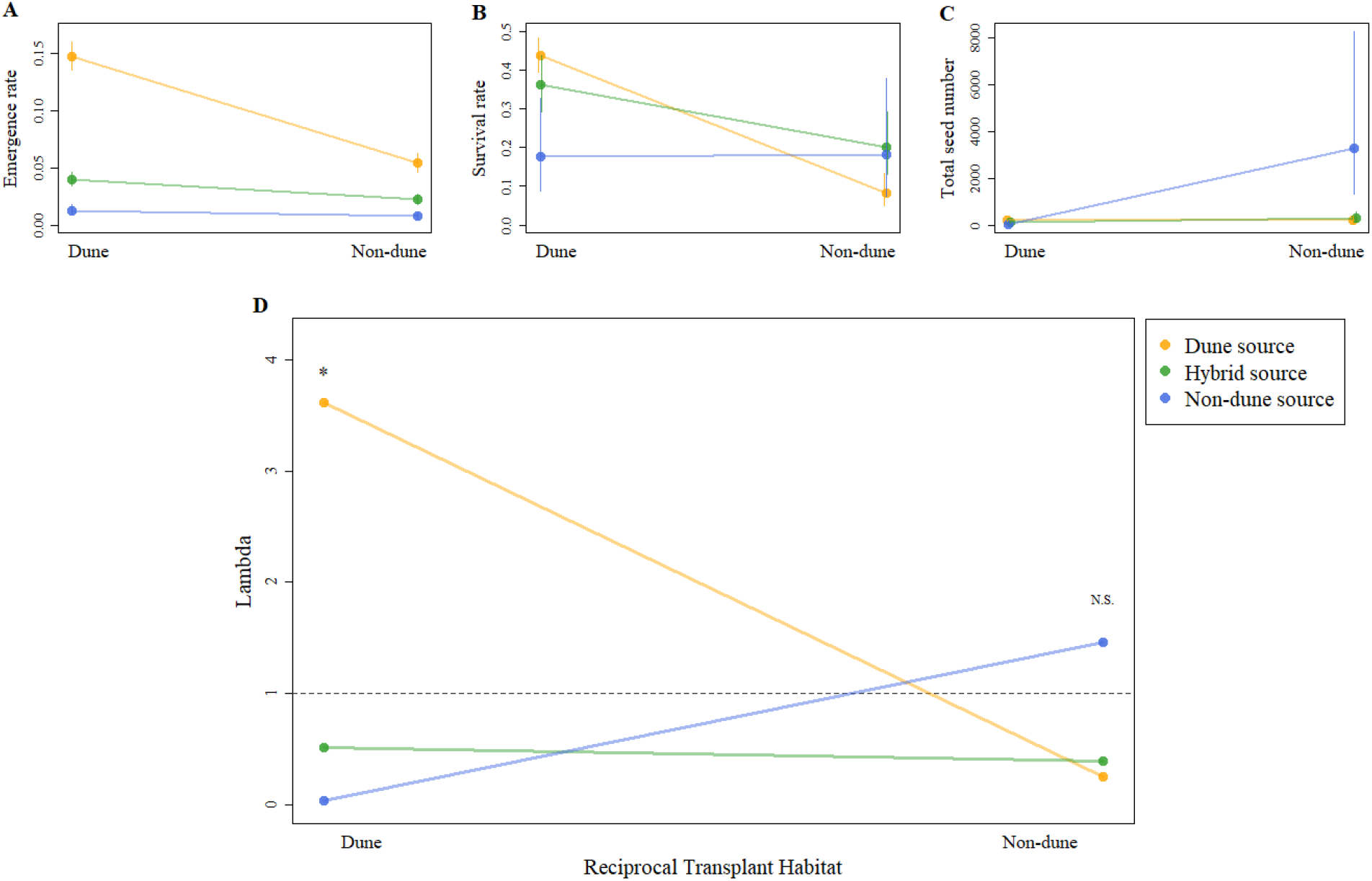
Reaction norms for fitness components (A-C) and population growth (D) for each source in both reciprocal transplant habitats. A) Mean emergence rate (number of seedlings that emerged per plot / number of seeds planted per plot) as estimated by GLM. B) Mean seedling-to-adult survival rate (number of plants that survived to flower per plot / number that emerged per plot) as estimated by GLM. C) Mean number of seeds produced (number of seeds counted from all collected heads, adjusted for the total number of recorded inflorescences and the proportion of seeds eaten prior to dispersal) as estimated by a negative binomial model. Error bars are 80% confidence intervals. Points and error bars and jittered horizontally to help visualization. D) Annual population growth rate (lambda) estimated by calculating the product of all estimated fitness components (emergence, survival, fecundity, seed survival). * and N.S. indicate p-values < and > 0.05, respectively, based on parametric bootstrapping (10,000 replicates) and 95% quantiles. The horizontal dotted line indicates lambda = 1; the boundary between population growth or decline.

While individual fitness components show a mix of effects with some evidence of local adaptation (fig 1A-C), the combined effects of all components generate lambda values that clearly indicate local adaptation (fig 1D). In both habitats, the local ecotype had greater fitness than foreign plants, with hybrids showing intermediate fitness, although this difference was only significant in the dune habitat (figs 1D, S2). Interestingly, our results also show that, in both habitats, only the local ecotype exhibited a lambda value >1, indicating that while each ecotype would be able to sustain populations in its native habitat, immigrants and hybrids would not.

LTRE analyses show that emergence contributed most to positive population growth of the local ecotype in the sand dunes, while fecundity was the most important component for positive growth of the local ecotype in the non-dune habitat (fig 2). Survival contributed the least to differences in lambda in either habitat (fig 2). These results indicate that different fitness components are responsible for positive population growth of local genotypes in each habitat, and that ecotype-specific strategies are likely contributing to these fitness components.

**Figure 2:**
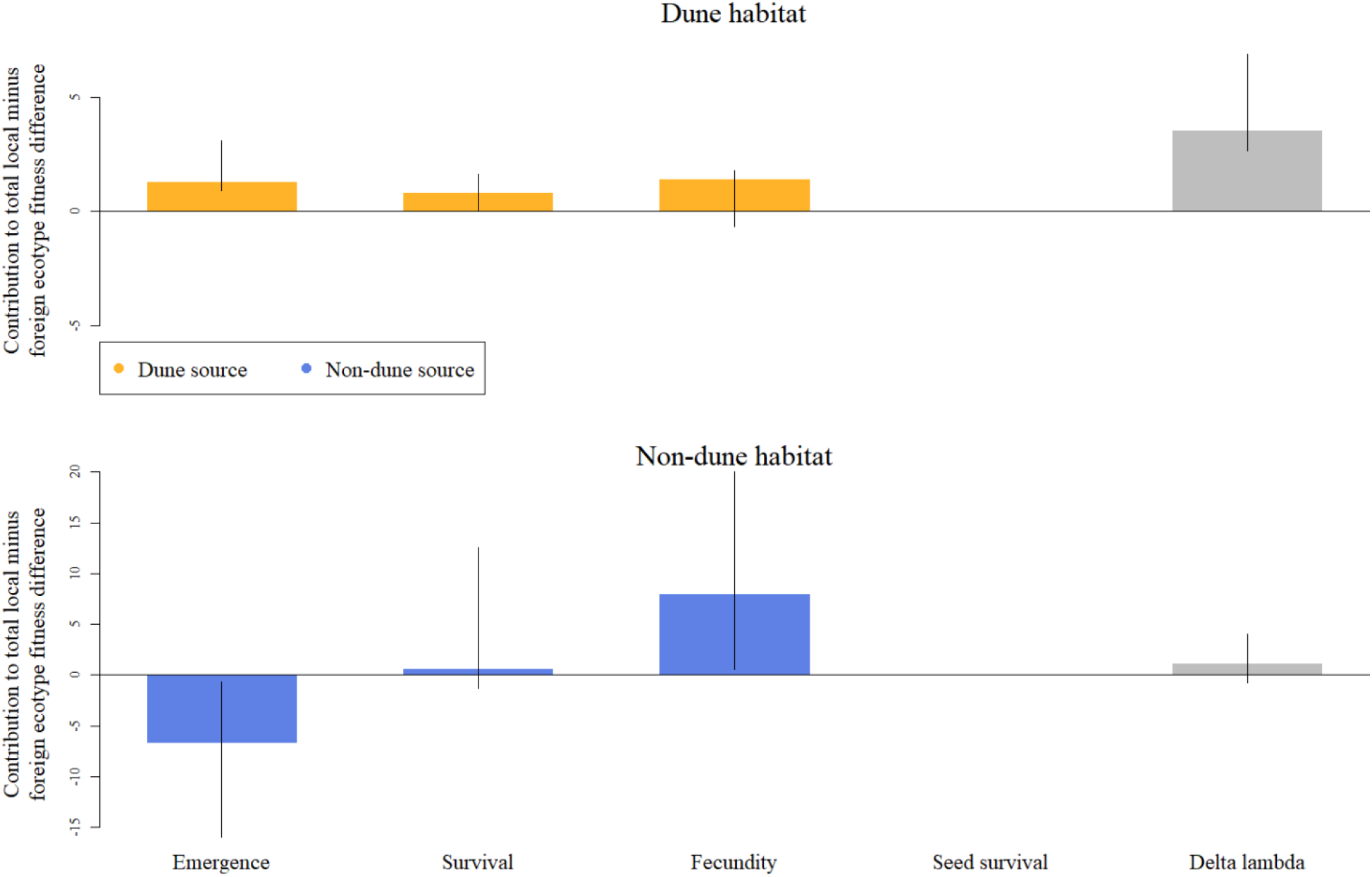
Results of a life table response experiment showing the relative difference in lambda estimates between ecotypes (local – foreign; grey bars) and the contribution of each fitness component to those differences in lambda (coloured bars) in each habitat; dune (top) and non-dune (bottom). Contribution values are quantified as the change in a given fitness component value (local – foreign) multiplied by the sensitivity of lambda to that given fitness component (note y-axis scales are different between panels). Error bars are 95% quantiles from parametric bootstrapping (10,000 replicates). Note that seed survival does not contribute to differences in lambda because we used a constant estimate for local and foreign individuals in each habitat.

### Allele frequency change due to selection at different life stages

After filtering we obtained genotype calls at 12,214 SNPs in 228 pre-selection plants of dune, non-dune, and hybrid source, and 103 plants that survived to maturity in either habitat (post-selection samples). This set of SNPs is assumed to contain neutral markers as well as SNPs physically or statistically linked to non-neutral loci.

We used principal component analysis (PCA) to determine the genetic structure of samples (figs S4, 3). PC1 explains 4.2% of the variation, while all other principal components individually explain less than 2% of the variation. This relatively low explanation of variation is typical when using thousands of SNPs (Gauch et al. 2019). As expected based on known population structure in this system (Andrew et al. 2013), samples largely separate by source along PC1, with hybrids intermediate to parents (fig S4). PC1 can therefore be interpreted as emphasizing the SNPs that are highly divergent between ecotypes and can be used to determine how genetically ‘dune-like’ versus ‘nondune-like’ samples are (fig S5).

When comparing plants that emerged and survived in the dune habitat to pre-selection samples, there was a genetic shift in all sources towards the dune side of PC1 (fig 3A). This suggests selection against foreign genotypes in the dune habitat. Interestingly, samples from all sources that survived in the non-dune habitat also shifted towards the dune-side of PC1 (fig 3B). While these latter shifts are smaller in magnitude and result from only a few surviving individuals, they suggest that dune alleles are favoured in both habitats at the emergence and survival stages. This result, while unexpected, makes sense considering the fitness component results where emergence was highest for dune plants in both habitats, even if the magnitude of this effect was greatest in the dunes (fig 1A). This suggests that these changes at early life stages are driven primarily by emergence, and not survival (figs 1A-B, 2).

**Figure 3:**
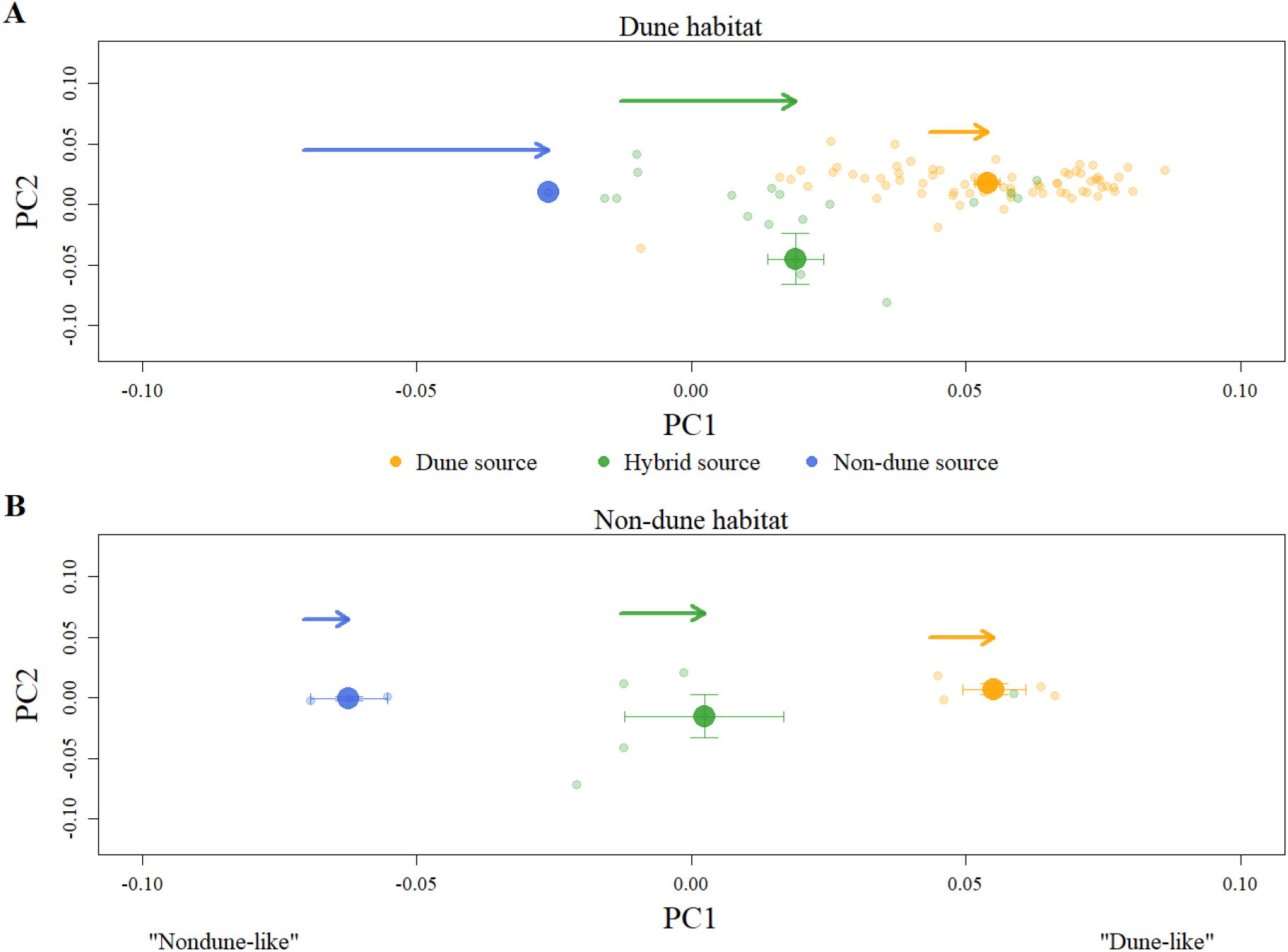
Principal component (PC) analysis based on post-selection samples genotyped at 12,214 SNPs from the A) dune habitat and the B) non-dune habitat. Scores for PC1 and PC2 for each sample (small points) as well as mean scores per source (large points) are plotted. Error bars represent standard error of the mean. Arrows indicate the direction and magnitude of shifts in mean PC1 scores between pre-(start of arrow, based on the means in figure S4) and post-selection (arrowhead) samples.

The shifts seen in figure 3 do not indicate specific genomic regions that are changing due to selection. To investigate this, we looked at the relationship between ecotypic differentiation and change in allele frequencies that occurred over early life stages. We found a significant positive relationship between ecotypic allele frequency difference (dune minus nondune) and allele frequency change (post-minus pre-selection) for hybrid samples in the dune habitat (r=0.14, p=0.004; fig 4). This indicates that the loci that changed the most in early life stages are often highly differentiated between ecotypes and that allele frequency change was generally in the expected direction (towards dune alleles); positive slope indicates that most loci had a higher frequency of dune alleles in post-selection samples (fig 4). We also found that loci exhibiting the largest shifts were distributed across the genome (figs S8, S9).

**Figure 4:**
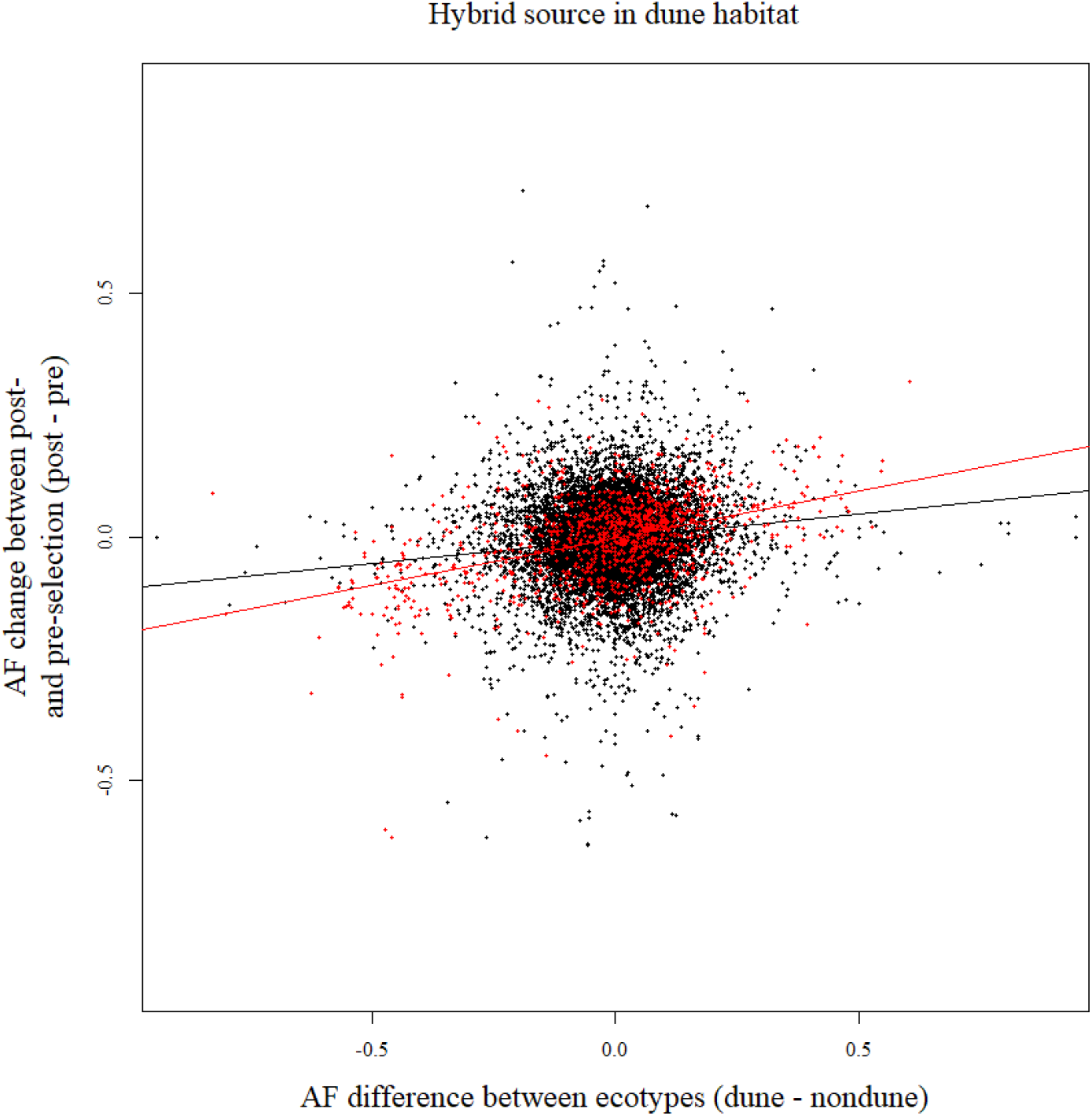
Relationship of allele frequency (AF) differences between ecotype (post-selection samples from non-experimental plants; dune minus non-dune) and allele frequency change during early life history (post-minus pre-selection) in the dune habitat in samples of hybrid source (r = 0.14, p = 0.004; all points). The relationship is stronger for inversion loci (r = 0.40, p = 0.003; red points) than for non-inversion loci (r = 0.085, p = 0.035; black points). P-values are based on one-sided tests using distributions from randomization tests (fig S7 A). Lines represent linear model fits (black = all points, red = inversion loci only).

We then specifically looked at how allele frequencies changed at loci inside inversions that are associated with ecotypic and habitat differentiation (Huang et al. 2020; Todesco et al. 2020). In plants of hybrid source, loci within inversions had a much stronger correlation (r=0.4, p=0.003) between ecotypic differentiation and allele frequency change than loci outside of inversions (r=0.085, p=0.035; fig 4). Interestingly, although inversion loci show more extreme patterns of allele frequency change overall, not every individual inversion is an outlier (fig S11, table S5). We also looked at allele frequency change in samples of dune source. We found the opposite pattern to that found in hybrid samples, where inversion loci in dunes samples generally had less allele frequency change than non-inversion loci (figs S6, S11, table S5). This finding is consistent with expectations of lower starting variation at adaptive loci in dune samples.

Finally, to investigate how alleles are affected at a later life stage, specifically via fecundity, we predicted the genotypes of seeds in the next generation by weighting individual PC scores by individual seed output (fig 5). There is a large shift towards non-dune alleles in hypothetical seeds produced by plants in the non-dune habitat (fig 5 dashed black arrow). This indicates that differences in seed output are linked to genotype differences, and that non-dune genotypes are correlated with higher seed production in their home environment. The smaller shift in predicted seeds in the dune habitat (fig 5 dashed grey arrow) likely reflects less differential fecundity (fig 1C), and potentially lower genotypic variation at loci associated with fecundity in this habitat. This result helps explain why, despite dune alleles being favoured in both habitats early in life, we observe divergent ecotypes in nature. We do not see dune alleles sweeping to fixation in both habitats because their early advantage is associated with lower seed production.

**Figure 5:**
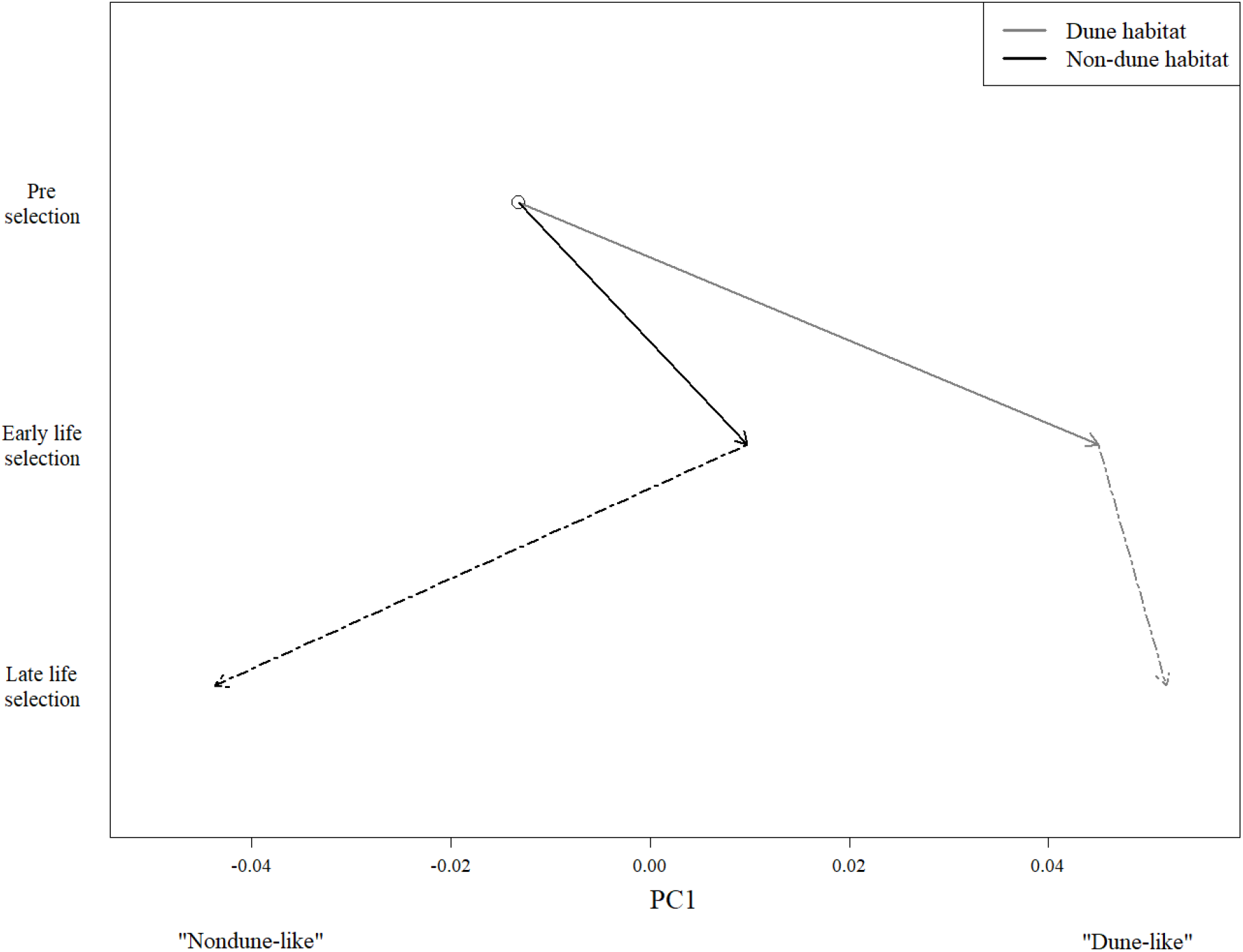
Shifts in mean PC1 scores following selection at different life stages in dune (grey) and non-dune (black) habitats. Mean PC1 scores are based on 12,214 SNPs from samples pooled across all sources (dune, hybrid, non-dune) that were grown under different conditions: greenhouse conditions (open circle, based on samples drawn in proportion to the number of each source planted in the field), the dune habitat (grey) and the non-dune habitat (black). Solid arrows indicate shifts following selection at early life stages and dashed arrows indicate shifts after adjusting for individual fecundity (i.e. after weighting individual PC1 scores by the number of seeds produced by each plant).

## Discussion

We investigated how selection on multiple fitness components occurring at different life history stages affects allele frequencies and contributes to absolute fitness of diverging ecotypes. It is well established that selection can vary across traits and life stages (Lande 1982), but it is less clear how this ultimately affects allele frequencies and persistence of diverging populations. By combining demographic population modelling and genomics, we show that selection on specific genomic regions and contrasting traits varies across life history stages to explain the maintenance of divergent adaptation.

### Fitness components and population growth

We found that the contributions of different fitness components to lambda were habitat dependent, and that this is likely caused by ecotype-specific life history strategies (figs 1, 2). Specifically, we saw that emergence was the most important contributor to increased fitness of the dune ecotype in the dunes, and fecundity was the most important contributor to increased success of the non-dune ecotype locally (fig 2). It has previously been reported in this system that dune plants produce larger seeds than non-dune plants (>2 times larger by mass) and that seed size is positively correlated with emergence and establishment (although seed size alone does not explain emergence patterns) (Ostevik et al. 2016). This suggests that large seeds that lead to higher emergence evolved in the dune habitat where these properties are important for population growth, possibly in response to frequent burial in shifting sand (Maun, 1998), or other early life history challenges. In contrast, high seed output, a trait observed in non-dune plants (fig 1C), positively impacts fecundity, which may be important for persistence in the non-dunes where unoccupied soil is less abundant (safe site limitation) and competition may be more acute (Andrew et al. 2012). The evolution of two contrasting life history strategies (i.e., seed quality vs quantity) in neighbouring, recently diverged ecotypes is a clear example of a trade-off driving divergence and incipient speciation (Ghalambor et al. 2004; Peterson et al. 2016).

Our lambda estimates also indicate local adaptation in this system (fig 1D). These results are qualitatively similar to previous analyses of the demographic data from this experiment using ASTER modeling (Ostevik et al. 2016). However, the current analysis allows inferences of absolute fitness as well as local adaptation.

While our finding that only the local ecotype has lambda >1 is realistic and supported by observations of distinct ecotypes in the field, there are several caveats to these estimates. First, all lambda values were calculated using a constant estimate of seed survival that was not measured in this system. We do not have data to determine the accuracy of our seed survival estimate, but a broad range of values (see methods) yielded qualitatively similar results. Another related consideration is that we used the same seed survival value for all sources in both habitats. We expect this assumption is, at least to some extent, wrong, as large dune seeds are likely to be predated more often than small non-dune seeds. However, if predation is high in the non-dunes where more vegetation and wildlife are found (Meiss et al. 2010), this may only strengthen our findings since dune seeds would have low seed survival in their foreign environment.

A second consideration when interpreting our results is inter-annual environmental variation. Observations of this system over multiple years reveal high variation in population size of the non-dune ecotype, which is suspected to be linked in part to precipitation (the dune ecotype appears to be more stable). The year of the experiment was drier than average, and we suspect that in dry years there is more water available to plants in the dunes due to pooling of water at reachable depths, relative to available water in the non-dunes where higher evaporation follows small precipitation events (i.e. the inverse texture hypothesis) (Noy-Meir 1973; Sala et al. 1988). This idea is supported by our findings of higher emergence and survival in the dune habitat, and observed low numbers of the natural non-dune ecotype in 2012 (pers. obs. by author).

While performance will vary from year to year, estimated differences in the importance of fitness components across habitats are likely robust. We found that while fecundity had the greatest contribution to lambda in the non-dunes, emergence was also important (fig 2). Therefore, in very dry years we expect that the negative contribution of emergence to the non-dune ecotype may be strong enough to cancel out the positive contribution of fecundity in its home environment (fig 2), possibly resulting in lambda values below 1. This may explain why in some years we see virtually no plants in the non-dune habitat (pers. obs. by author); a seed bank here may facilitate large population size fluctuations (Cohen 1966). In contrast, in wet years when emergence is higher in the non-dunes, we expect that some dune individuals may succeed in their foreign habitat, in addition to an abundance of local individuals (boom years have been observed but less often than non-boom years). This speculation supports previous reports from genetic analysis of asymmetric gene flow, where rates of gene flow are higher from dunes to non-dunes rather than vice versa (Andrew et al. 2013). This suggests that the derived, dune ecotype may have the potential to inhabit a broader range of habitats than the ancestral type.

### Allele frequency change due to selection at different life stages

We demonstrated that selection at different life history stages affects allele frequencies, and that the relative magnitude of these effects are habitat dependent (fig 5). When assessing allele frequency shifts following emergence and survival, we saw the largest shifts in the dune habitat (fig 3), suggesting stronger selection in this habitat at early stages. This makes sense in light of our findings that emergence is the most important fitness component for success in the dune habitat (figs 1, 2). The most dramatic shift in the dunes was for non-dune samples (fig 3A); while only one plant survived, its genotype was considerably more ‘dune-like’ than the average of pre-selection samples. The low survival as well as the genetic composition of the sole surviving non-dune plant supports our conclusion that foreign alleles are selected against in the dune habitat at early life stages. The smaller shift seen for the dune samples likely indicates low amounts of variation for selection to act on, particularly at loci of adaptive interest. The hybrid samples show a moderate shift towards the dune side of PC1. While the hybrids should have the most genetic variation and thus the largest potential for change, these individuals may be at a disadvantage if heterozygosity at adaptive loci, or new combinations of alleles, are selected against. This is supported by our lambda estimates that show hybrids having low fitness, (fig 1D), suggesting that heterozygosity or disruption of adaptive allele combinations are disadvantageous. For all sources, a similar pattern is seen in the non-dune habitat, but shifts are generally lower in magnitude (fig 3B). Our PCA plots show how allele frequencies shift towards dune alleles at early stages in both habitats, however, they do not inform on what specific genomic regions are responsible for these shifts.

When examining how specific loci changed during the experiment, we found that loci with large allele frequency shifts were distributed across the genome (figs S8, S9). The distribution of allele frequency change differed by source, highlighting differences in starting variation and linkage disequilibrium in plants of hybrid versus dune source, as well as differences that are expected due to drift (figs S8, S9). When focusing on samples of hybrid source, which had higher starting variation, we found a significant positive correlation between allele frequency change during early life and genomic differentiation between ecotypes, suggesting an increase in dune alleles at potentially adaptive loci (fig 4). However, we also see some highly differentiated loci that did not change in frequency during the experiment, as well as a number of loci with large allele frequency change at non-differentiated loci (fig 4). The former group likely includes loci that are important at life stages not measured here, and the latter group is likely the result of random drift or selection acting in year-specific ways that might not promote divergence.

Seven large putative inversions have been found to be differentiated between ecotypes in this system and proposed to contain adaptive alleles (Huang et al. 2020; Todesco et al. 2020). We found support for three of these inversions to be important during early life stages. Inversion pet17.01 showed significantly (p<0.05) increased allele frequency change in the hybrid source and decreased allele frequency change in the dune source (table S5). Inversions pet05.01 and pet11.01 also showed patterns consistent with our expectations, but these patterns were not always significant (table S5). These results support that strong barriers to gene flow are acting at loci in these inversions such that dune samples are largely fixed for dune alleles (Huang et al. 2020), and that ongoing selection maintains existing allelic composition. Only the artificially generated hybrids had the necessary allelic variation for selection to be observed. These three inversions have been found to be associated with seed size (pet11.01; Todesco et al. 2020) and/ or soil and vegetative cover characteristics (pet05.01, pet11.01, pet17.01; Huang et al. 2020). While three of the remaining inversions also have significant associations with seed size (pet09.01, pet14.01; Todesco et al. 2020) or habitat differences (pet07.01; Huang et al. 2020), our analyses show mixed support for selection on these inversions during early life stages (table S5). Perhaps these inversions are subject to selection at different life history stages or during environmental conditions that were not present in the year of the experiment. Also, our dataset may not have ample resolution to detect a strong trend in some inversions (eg. pet14.01 lacks enough starting variation for conclusions to be drawn).

Loci driving the observed allele frequency shifts are likely not the direct targets of selection; GBS data provide sequence for only a few thousand representative sites throughout the genome. Additionally, our approach of looking at allele frequency change will predominantly be relevant for loci with additive gene action. Despite these caveats, we found that allele frequency change during early life was largely driven by loci in several segregating inversions (fig 4, table S5). This suggests that in a single generation we see allele frequency change due to divergent adaptation that explains observed allele frequencies at many loci. These results provide a basis for future investigations of functional elements located in identified inversions that are important for adaptation during early life history.

Finally, when assessing predicted allele frequency shifts following fecundity, we saw the largest shifts in the non-dune habitat (fig 5). This indicates that fecundity affects allele frequencies, and corroborates our earlier finding that fecundity contributes most to positive population growth in the non-dune habitat (fig 2). While this result is based on data from only a few non-dune individuals that survived during the experiment, data collected in other years supports findings of high fecundity in this ecotype (Ostevik et al. 2016). In the dune habitat, we see a smaller shift in allele frequencies after adjusting for fecundity (fig 5). Presumably, this small shift is due to lower genetic variation at loci linked to fecundity. Note that our predictions of allele frequency change in the next generation assume that allele contributions from pollen are the same as contributions from the maternal plant. While this assumption is surely violated, we think that the male and female genetic contributions within an ecotype are similar enough to make this analysis useful.

## Conclusions

In summary, using a combined set of analyses allowed us to obtain a clearer picture of the gene flow-selection balance in this system. We identified different life history strategies in neighbouring ecotypes that appear to have evolved by optimization of different fitness components, which in turn contribute to the persistence of each ecotype in its respective habitat. We also showed that the maintenance of divergent adaptation in this system is mediated via habitat and life stage-specific selection that alters allele frequencies. Lastly, we were able to observe allele frequency shifts during early life stages at specific loci that have previously been shown to be under selection (Huang et al. 2020; Todesco et al. 2020). These findings demonstrate that ecological selection can be complex and environmentally dependent, and can act on different genomic regions at different life history stages. Our work in this system provides an unusual investigation into the details of how selection operates in nature and contributes to our fundamental understanding of evolution. Additionally, our findings can be applied to understanding how populations adapt to new habitats, how small populations persist despite on-going gene flow, and to the conservation of diversity more generally.

## Acknowledgments

We thank Great Sand Dunes National Park and Preserve, A. Valdez, P. Bovin, F. Bunch, J. Herndon, S. Tittes, C. Smith, M. Peterson, and J. Stanley. Thank you to A. Kuzmin and the UBC Biodiversity Research Center for help with sequencing. This work was supported by NSERC Postgraduate Scholarships (AMG and KLO), NSF IGERT grant number 1144807 (AMG), NSERC Discovery Grant number 327475 (LHR).

## Data Accessibility

Demographic data was obtained from 10.5061/dryad.223p4. Genetic sequence data has been deposited on the Sequence Read Archive (SRA) and will be available upon publication at www.ncbi.nlm.nih.gov/sra/PRJNA623572.

## Author contributions

KLO and LHR conceived of the reciprocal transplant experiment and associated sequencing. KLO performed the reciprocal transplant experiment and associated demographic and DNA sequence data collection. AMG performed analyses. All authors contributed to interpretation of results and writing of the manuscript.

## Notes

### Competing Interest Statement

The authors have declared no competing interest.

## References

Alexander, H. M., Cummings, C. L., Kahn, L., & Snow, A. A. (2001). Size Variation and Predation of Seeds Produced by Wild and Crop–Wild Sunflowers. American Journal of Botany, 88(4), 623–627. https://doi.org/10.2307/2657061

Anderson, J., Lee, C.-R., & Mitchell-Olds, T. (2014). Strong Selection Genome-wide Enhances Fitness Tradeoffs Across Environments and Episodes of Selection. Evolution, 68(1), 16–31. https://doi.org/10.1111/evo.12259

Andrew, R. L., Kane, N. C., Baute, G. J., Grassa, C. J., & Rieseberg, L. H. (2013). Recent nonhybrid origin of sunflower ecotypes in a novel habitat. Molecular Ecology, 22(3), 799–813. https://doi.org/10.1111/mec.12038

Andrew, R. L., Ostevik, K. L., Ebert, D. P., & Rieseberg, L. H. (2012). Adaptation with gene flow across the landscape in a dune sunflower. Molecular Ecology, 21(9), 2078–2091. https://doi.org/10.1111/j.1365-294X.2012.05454.x

Andrew, R. L., & Rieseberg, L. H. (2013). Divergence is focused on few genomic regions early in speciation: Incipient speciation of sunflower ecotypes. Evolution, 67(9), 2468–2482. https://doi.org/10.1111/evo.12106

Andrews, S. (2010). FastQC: A quality control tool for high throughput sequence data. Retrieved from http://www.bioinformatics.babraham.ac.uk/projects/fastqc

Antonovics, J. (2006). Evolution in closely adjacent plant populations X: Long-term persistence of prereproductive isolation at a mine boundary. Heredity, 97(1), 33–37. https://doi.org/10.1038/sj.hdy.6800835

Badouin, H., Gouzy, J., Grassa, C. J., Murat, F., Staton, S. E., Cottret, L., … Langlade, N. B. (2017). The sunflower genome provides insights into oil metabolism, flowering and Asterid evolution. Nature, 546, 148–152. https://doi.org/10.1038/nature22380

Brady, K. U., Kruckeberg, A. R., Bradshaw Jr., H. D. (2005). Evolutionary ecology of plant adaptation to serpentine soils. Annual Review of Ecology, Evolution, and Systematics, 36, 243–266.

Bolger, A. M., Lohse, M., & Usadel, B. (2014). Trimmomatic: A flexible trimmer for Illumina sequence data. Bioinformatics, 30(15), 2114–2120. https://doi.org/10.1093/bioinformatics/btu170

Caisse, M., & Antonovics, J. (1978). Evolution in closely adjacent plant populations: IX. Evolution of reproductive isolation in clinal populations. Heredity, 40(3), 371–384. https://doi.org/10.1038/hdy.1978.44

Campbell, D. R., & Waser, N. M. (2007). Evolutionary dynamics of an Ipomopsis hybrid Zone: Confronting models with lifetime fitness data. The American Naturalist, 169(3), 298–310.

Caswell, H. (2001). Matrix Population Models: Construction, Analysis, and Interpretation. Sinauer Associates.

Cohen, D. (1966). Optimizing Reproduction in a Randomly Varying Environment. Journal of Theoretical Biology, 12, 119–129.

Cotto, O., Sandell, L., Chevin, L., & Ronce, O. (2019). Maladaptive Shifts in Life History in a Changing Environment. The American Naturalist, 194(2), 558–573. https://doi.org/10.1086/702716

Coulson, T., Kruuk, L. E. B., Tavecchia, G., Pemberton, J. M., & Clutton-Brock, T. H. (2003). Estimating Selection on Neonatal Traits in Red Deer Using Elasticity Path Analysis. Evolution, 57(12), 2879–2892.

Danecek, P., Auton, A., Abecasis, G., Albers, C. A., Banks, E., DePristo, M. A., … Group, 1000 Genomes Project Analysis. (2011). The variant call format and VCFtools. Bioinformatics, 27(15), 2156–2158. https://doi.org/10.1093/bioinformatics/btr330

Dechaine, J. M., Burger, J. C., & Burke, J. M. (2010). Ecological patterns and genetic analysis of post-dispersal seed predation in sunflower (Helianthus annuus) crop-wild hybrids. Molecular Ecology, 19(16), 3477–3488. https://doi.org/10.1111/j.1365-294X.2010.04740.x

Doyle, J. J., & Doyle, J. L. (1987). A rapid DNA isolation procedure for small quantities of fresh leaf tissue. Phyochemical Bulletin, 19(1), 11–15.

Egan, S. P., Ragland, G. J., Assour, L., Powell, T. H. Q., Hood, G. R., Emrich, S., … Feder, J. L. (2015). Experimental evidence of genome-wide impact of ecological selection during early stages of speciation-with-gene-flow. Ecology Letters, 18(June), 817–825. https://doi.org/10.1111/ele.12460

Ehrlen, J., & Munzbergova, Z. (2009). Timing of Flowering: Opposed Selection on Different Fitness Components and Trait Covariation. The American Naturalist, 173(6), 819–830. https://doi.org/10.1086/598492

Felsenstein, J. (1981). Skepticism Towards Santa Rosalia, or Why are There so Few Kinds of Animals? Evolution, 35(1), 124–138.

Ferris, K. G., & Willis, J. H. (2018). Differential adaptation to a harsh granite outcrop habitat between sympatric Mimulus species. Evolution, 72, 1225–1241. https://doi.org/10.1111/evo.13476

Gauch, H. G. J., Qian, S., Piepho, H.-P., Zhou, L., & Chen, R. (2019). Consequences of PCA graphs, SNP codings, and PCA variants for elucidating population structure. PLoS ONE, 1–26.

Griffith, A. B. (2010). Positive effects of native shrubs in *Bromus tectorum* demography. Ecology, 91(1), 141–154.

Ghalambor, C. K., Reznick, D. N., & Walker, J. A. (2004). Constraints on Adaptive Evolution: The Functional Trade-Off between Reproduction and Fast-Start Swimming Performance in the Trinidadian Guppy (Poecilia reticulata). The American Naturalist, 164(1), 38–50.

Gompert, Z., Comeault, A. A., Farkas, T. E., Feder, J. L., Parchman, T. L., Buerkle, C. A., & Nosil, P. (2014). Experimental evidence for ecological selection on genome variation in the wild. Ecology Letters, 17(3), 369–379. https://doi.org/10.1111/ele.12238

Harrison, S., & Rajakaruna, N. (2011). Serpentine: The evolution and ecology of a model system. Berkeley: University of California Press.

Hoban, S., Kelley, J. L., Lotterhos, K. E., Antolin, M. F., Bradburd, G., Lowry, D. B., … Whitlock, M. C. (2016). Finding the Genomic Basis of Local Adaptation: Pitfalls, Practical Solutions, and Future Directions. The American Naturalist, 188(4), 379–397. https://doi.org/10.1086/688018

Horvitz, C. C., Coulson, T., Tuljapukar, S., & Schemske, D. W. (2010). A new way to integrate selection when both demography and selection gradients vary over time. International Journal of Plant Sciences, 171(9), 945–959. https://doi.org/10.1086/657141.A

Huang, K., Andrew, R. L., Owens, G. L., Ostevik, K. L., & Rieseberg. (2020). Multiple chromosomal inversions contribute to adaptive divergence of a dune sunflower ecotype. Molecular Ecology, 29, 2535–2549.

Jain, S. K., & Bradshaw, A. D. (1966). Evolutionary divergence among adjacent plant populations. I. The evidence and its theoretical analysis. Heredity, 21, 407–441.

Kawecki, T., & Ebert, D. (2004). Conceptual issues in local adaptation. Ecology Letters, 7, 1225–1241. https://doi.org/10.1111/j.1461-0248.2004.00684.x

Kruckeberg, A. R. (1950). An experimental inquiry into the nature of endemism on serpentine soils. University of California Berkeley.

Lande, R. (1982). A Quantitative Genetic Theory of Life History Evolution Russell Lande. Ecology, 63(3), 607–615.

Li, H. (2011). A statistical framework for SNP calling, mutation discovery, association mapping and population genetical parameter estimation from sequencing data. Bioinformatics, 27(21), 2987–2993. https://doi.org/10.1093/bioinformatics/btr509

Li, H. (2013). Aligning sequence reads, clone sequences and assembly contigs with BWA-MEM. ArXiv, 0(00), 1–3. https://doi.org/10.1186/s13756-018-0352-y

Luu, K., Bazin, E., & Blum, M. G. B. (2017). PCAdapt: An R package to perform genome scans for selection based on principal component analysis. Molecular Ecology Resources, 17, 67–77. https://doi.org/10.1111/1755-0998.12592

Maun, M. A. (1998). Adaptations of plants to burial in coastal sand dunes. Canadian Journal of Botany, 76, 713–738.

Maynard Smith, J. (1966). Sympatric Speciation. American Naturalist, 100, 637–650. https://doi.org/10.1086/285850

McNeilly, T., & Antonovics, J. (1968). Evolution in closely adjacent plant populations. IV. Barriers to gene flow. Heredity, 23, 205–218.

Meiss, H., Le Lagadec, L., Munier-Jolain, N., Waldhardt, R., & Petit, S. (2010). Weed seed predation increases with vegetation cover in perennial forage crops. Agriculture, Ecosystems and Environment, 138, 10–16. https://doi.org/10.1016/j.agee.2010.03.009

Millard, S. P. (2013). EnvStats: An R package for environmental statistics. New York: Springer.

Morris, W. F., & Doak, D. F. (2002). Quantitative Conservation Biology: Theory and Practice of Population Viability Analysis. Sinauer Associates.

Najoshi. (2017). Sabre. Najoshi. Retrieved from https://github.com/najoshi/sabre

Nosil, P., Crespi, B. J., & Sandoval, C. P. (2002). Host-plant adaptation drives the parallel evolution of reproductive isolation. Nature, 417, 440–443.

Noy-Meir, I. (1973). Desert Ecosystems: Environment and Producers. Review of Ecology and Systematics, 4, 25–51.

Ostevik, K. L., Andrew, R. L., Otto, S. P., & Rieseberg, L. H. (2016). Multiple reproductive barriers separate recently diverged sunflower ecotypes. Evolution, 70(10), 2322–2335. https://doi.org/10.1111/evo.13027

Peterson, M. L., Kay, K. M., & Angert, A. L. (2016). The scale of local adaptation in Mimulus guttatus: Comparing life history races, ecotypes, and populations. New Phytologist, 211, 345–356. https://doi.org/10.1111/nph.13971

Poland, J. A., Brown, P. J., Sorrells, M. E., & Jannink, J. L. (2012). Development of high-density genetic maps for barley and wheat using a novel two-enzyme genotyping-by-sequencing approach. PLoS ONE, 7(2). https://doi.org/10.1371/journal.pone.0032253

R Core Team. (2018). R: A language and environment for statistical computing. R Foundation for Statistical Computing, Vienna, Austria. https://www.R-project.org/

Rohland, N., & Reich, D. (2012). Cost-effective, high-throughput DNA sequencing libraries for multiplexed target capture. Genome Research, 22, 939–946. https://doi.org/10.1101/gr.128124.111.22

Sala, O. E., Parton, W. J., Joyce, L. A., & Lauenroth, W. K. (1988). Primary Production of the Central Grassland Region of the United States. Ecology, 69(1), 40–45.

Sambatti, J. B. M., & Rice, K. J. (2006). Local Adaptation, Patterns of Selection, and Gene Flow in the Californian Serpentine Sunflower (Helianthus exilis). Evolution, 60(4), 696–710.

Schluter, D. (1993). Adaptive Radiation in Sticklebacks: Size, Shape, and Habitat Use Efficiency. Ecology, 74(3), 699–709.

Slatkin, M. (1987). Gene flow and the geographic structure of natural populations. Science, 236, 787–792.

Smith, M., Caswell, H., & Mettler-Cherry, P. (2005). Stochastic Flood and Precipitation Regimes and the Population Dynamics of a Threatened Floodplain Plant. Ecological Applications, 15(3), 1036–1052.

Takada, T., & Shefferson, R. (2018). The long and winding road of evolutionary demography: preface. Population Ecology, 60, 3–7. https://doi.org/10.1007/s10144-018-0622-9

Thurman, T. J., & Barrett, R. D. H. (2016). The genetic consequences of selection in natural populations. Molecular Ecology, 25, 1429–1448. https://doi.org/10.1111/mec.13559

Tigano, A., & Friesen, V. L. (2016). Genomics of local adaptation with gene flow. Molecular Ecology, 25, 2144–2164. https://doi.org/10.1111/mec.13606

Todesco, M., Owens, G. L., Bercovich, N., Légaré, J., Soudi, S., Burge, D. O., … Rieseberg, L. H. (2020). Massive haplotypes underlie ecotypic differentiation in sunflowers. Nature.

van Noordwijk, A. J., & de Jong, G. (1986). Acquisition and Allocation of Resources: Their Influence on Variation in Life History Tactics. The American Naturalist, 128(1), 137–142. https://doi.org/10.1086/284547

Venable, D. L. (1992). Size-Number Trade-Offs and the Variation of Seed Size with Plant Resource Status. The American Naturalist, 140(2), 287–304.

Wu, C. (2001). The genic view of the process of speciation. Journal of Evolutionary Biology, 14, 851–865.

Zhulidov, P. A., Bogdanova, E. A., Shcheglov, A. S., Vagner, L. L., Khaspekov, G. L., Kozhemyako, V. B., … Shagin, D. A. (2004). Simple cDNA normalization using kamchatka crab duplex-specific nuclease. Nucleic Acids Research, 32(3). https://doi.org/10.1093/nar/gnh031

