## Supplemental Materials for "Contrasting selection at multiple life stages maintains divergent adaptation between sunflower ecotypes"

### Supplemental Methods

*Non-experimental plant collection*

To obtain genotypes for each ecotype, we used adult plants from naturally occurring (non-experimental) populations. Samples from non-experimental plants were collected from 27 dune individuals from one of the source populations (1701: 37.803N, 105.524W) and 57 non-dune individuals from two of the source populations (1791: 37.813N, 105.515W, 2001: 37.757N, 105.507W).

*Genotyping-by-sequencing library preparation*

Genomic DNA samples with low yield or poor absorbance ratios (260/280 nm or 260/230 nm) were re-extracted. Genotyping-by-sequencing (GBS) library preparation was performed according to a modified version of the Poland et al. (2012) protocol. Briefly, 100 ng of genomic DNA was digested with *Pst*I-HF and *Msp*I. Illumina adapters with PCR primer sites and unique 4-8 base pair (bp) barcodes were ligated to digested fragments. Up to 192 barcoded samples were pooled and concentrated using SeraMag Speed Beads made in-house (Rohland and Reich 2012). PCR using 12 cycles was performed to amplify fragments. To reduce the amount of high-copy fragments, libraries were denatured, allowed to re-hybridize, digested with Duplex Specific Nuclease (Zhulidov et al. 2004), and then undigested fragments were amplified with an additional 12 cycles of PCR (Todesco et al. 2020). Size selection for 300-800 bp fragments was done using gel electrophoresis on a 1.5% agarose gel; DNA was extracted using a Zymoclean Gel DNA Recovery kit (Zymo Research, Irvine, USA). Quality of libraries was accessed using an Agilent 2100 Bioanalyzer (Agilent Technologies, Santa Clara, CA, USA), and 100 bp read paired-end sequencing was performed on 3 lanes of an Illumina HiSeq 2000 (Illumina Inc., San Diego, CA, USA).

*Raw sequence data processing*

Raw short read data were demultiplexed using Sabre (github.com/najoshi/sabre, accessed August 2017) and quality checked using FastQC (Andrews 2010). Reads were filtered and trimmed using Trimmomatic version 0.36 (Bolger et al. 2014) as follows: Adapters were removed using the command ILLUMINACLIP:TruSeq3-PE-2.fa:2:30:4, restriction sites were removed, any bases with quality below 15 were removed from the leading and trailing ends of reads, reads were removed if total average quality or average quality in a 4 bp window was below 15, reads of less than 75 bp were removed.

*Random sampling of pre-selection hybrid types*

The amount of seed generated from BCN hybrids (back cross hybrids generated with non-dune pollen and equal proportions of dune and non-dune cytoplasms) was lower than the other hybrid types. This lower seed number prevented the BCN hybrid type from being grown in greenhouse conditions to serve as pre-selection genotypes. To account for the lack of BCN hybrids in pre-selection samples, we randomly sampled individuals of non-dune and F1 source to represent the missing genotypes. Samples were randomly selected in the following proportions: 0.5 non-dune, 0.25 F1D, 0.25 F1N. To make up the final hybrid pre-selection pool, samples of the other hybrid types were also randomly sampled to obtain proportions equal to those planted in the field. All random sampling was performed 1000 times and mean values (of either PC1 scores or allele frequencies) and 95% confidence intervals (of PC1 scores) across all repetitions were calculated.

*Estimating genotypes when treating each inversion as a single locus*

To test the effects of selection on putative chromosomal inversions, we followed the method described in Huang et al. 2020 for classifying each inversion as a single locus. Briefly, using SNPs for each inversion separately we calculated PCAs using the R package PCAdapt (Luu et al. 2017) version 4.1.0 with K=5 and default arguments. Next, PC1 scores were clustered using the R function kmeans in the R package stats v3.6.2. K-means cluster assignment was then used to assign each sample a genotype for each inversion. Clusters associated with the non-dune side of the PC1 axis were always assigned the reference genotype, while the dune associated cluster was assigned the alternate genotype. Finally, genotype calls were used to calculate allele frequencies as described in the main text. We excluded inversion pet09.02 from this analysis since we could not confidently assign individuals a genotype due to lack of distinct clustering.

### Supplemental tables

Table S1: Number of samples of each source from greenhouse (pre-selection) and post-selection field habitats that survived to reproduce, were successfully sequenced (yielded sufficient data) and included in genetic analyses.

| | Pre-selection | Dune habitat | Non-dune habitat |
|---|---|---|---|
| Dune source | 56 | 70 | 4 |
| Hybrid source | 81 | 21 | 5 |
| Non-dune source | 91 | 1 | 2 |

Table S2: Comparison of simple versus mixed model for modeling fitness components. Correlation ($R^2$) and corresponding p-values for the linear relationship between coefficients from simple (Fitness component ~ Source * Habitat) and mixed (Fitness component ~ Source * Habitat + (1|Population) + (1|Plot)) models of each fitness component. Emergence and survival data were fit with a binomial generalized linear model or a generalized linear mixed model. Seed number was fit with a negative binomial model or negative binomial mixed model. The coefficients from simple and mixed models are similar; we present results from the simple models in the main text.

| Fitness component modelled | $R^2$ | P |
|---|---|---|
| Emergence | 0.9956 | 4.572e-06 |
| Survival | 0.9334 | 0.001083 |
| Seed number | 0.9567 | 0.0004558 |

Table S3: Cutoff values used in iterative filtering of missing data. Filters applied consecutively from row 1-6. Number of remaining samples and SNPs after each filtering step are shown. The starting sample and SNP values were 437 and 680,100, respectively.

| | Filter | Number of samples remaining | Number of SNPs remaining |
|---|---|---|---|
| 1 | Removed samples missing 99.99% of sites | 431 | 680,100 |
| 2 | Removed sites that are in <20% of samples | 431 | 47,353 |
| 3 | Removed samples missing >94.8% of sites | 419 | 47,353 |
| 4 | Removed sites that are in <50% of samples | 419 | 25,797 |
| 5 | Removed samples missing >70% of sites | 411 | 25,797 |
| 6 | Removed sites that are in <75% of samples | 411 | 12,214 |

Table S4 A: Model statistics generated by anova type II test for binomial generalized linear model of seedling emergence rates (Emergence ~ Source * Habitat). Df: degrees of freedom.

| | Chi-square | Df | P |
|---|---|---|---|
| Source | 271.587 | 2 | < 2e-16 |
| Habitat | 72.756 | 1 | < 2e-16 |
| Source * Habitat | 5.458 | 2 | 0.06528 |

Table S4 B: Model statistics generated by anova type II test for binomial generalized linear model of seedling to adult survival rates (Survival ~ Source * Habitat). Df: degrees of freedom.

| | Chi-square | Df | P |
|---|---|---|---|
| Source | 2.395 | 2 | 0.30193 |
| Habitat | 31.924 | 1 | 1.603e-08 |
| Source * Habitat | 6.574 | 2 | 0.03737 |

Table S4 C: Model statistics generated by anova type II test for negative binomial model of seed output (Seed number ~ Source * Habitat). Df: degrees of freedom.

| | Chi-square | Df | P |
|---|---|---|---|
| Source | 9.9992 | 2 | 0.006741 |
| Habitat | 3.4967 | 1 | 0.061490 |
| Source * Habitat | 7.2374 | 2 | 0.026817 |

Table S5: Quantifying allele frequency (AF) change during early life stages in each inversion. Allele frequencies for pre- and post-selection samples of each source (dune and hybrid) were calculated for each inversion (inversions treated as single loci; pet09.02 not included due to inconclusive genotype assignment). Differences in proportions (AF x n x 2) of each allele (reference and alternate) between pre- and post-selection were determined using a Fisher's exact test (FET). AF change was calculated as the difference between pre- and post-selection AFs (post minus pre). The magnitude of change was compared to the distribution of AF change for all non-inversion loci; percentiles are reported. Additionally, the distribution of AF change in inversions (all SNPs treated separately) was compared to the distribution of AF change in all non-inversion loci using a two-sample linear rank test (+ or – indicates whether a significant change was greater or less than average).

| **Sample source** | **Inversion code** | **AF pre-selection** | **AF post-selection** | **p-value (FET)** | **AF change** | **Magnitude** | **p-value (rank-test)** |
|---|---|---|---|---|---|---|---|
| Dune | pet05.01 | 0.91 | 0.91 | 1 | -0.0036 | 6th | < 0.001 (-) |
| Dune | pet07.01 | 0.96 | 0.97 | 0.52 | 0.016 | 24th | 0.001 (-) |
| Dune | pet09.01 | 0.84 | 0.79 | 0.33 | -0.054 | 66th | 0.002 (-) |
| Dune | pet09.02 | NA | NA | NA | NA | NA | 0.33 |
| Dune | pet11.01 | 1 | 0.99 | 1 | -0.0071 | 11th | < 0.001 (-) |
| Dune | pet14.01 | 0.18 | 0.21 | 0.63 | 0.029 | 41st | 0.79 |
| Dune | pet17.01 | 0.98 | 0.98 | 1 | -0.0036 | 6th | < 0.001 (-) |
| Hybrid | pet05.01 | 0.57 | 0.71 | 0.11 | 0.14 | 96th | 0.039 (+) |
| Hybrid | pet07.01 | 0.78 | 0.69 | 0.23 | -0.092 | 87th | 0.012 (+) |
| Hybrid | pet09.01 | 0.61 | 0.69 | 0.38 | 0.077 | 81st | 0.002 (+) |
| Hybrid | pet09.02 | NA | NA | NA | NA | NA | 0.017 (-) |
| Hybrid | pet11.01 | 0.63 | 0.79 | 0.07 | 0.15 | 97th | 0.001 (+) |
| Hybrid | pet14.01 | 0.039 | 0.071 | 0.40 | 0.032 | 45th | 0.99 |
| Hybrid | pet17.01 | 0.77 | 0.95 | 0.007 | 0.18 | 98th | 0.002 (+) |

**Supplemental figures**

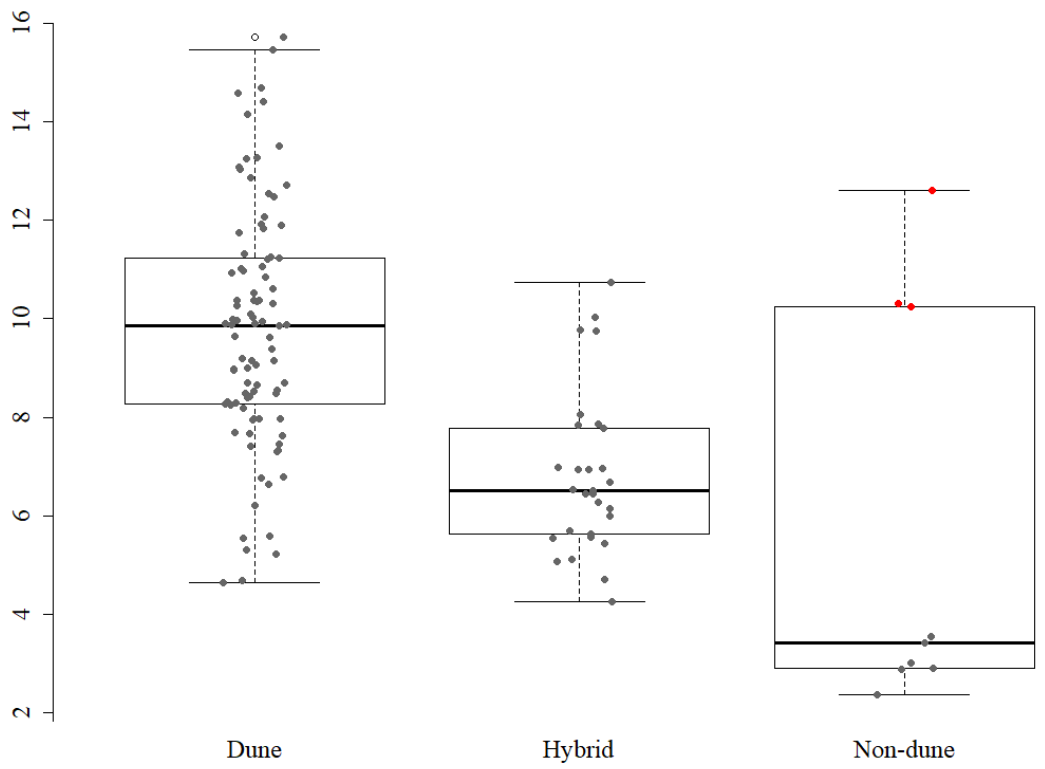

Figure S1: Boxplot of seed mass from seeds produced by plants of each source in the reciprocal transplant experiment. Plants that survived in their foreign habitat and produced seeds outside of their ecotypic seed weight range and inside the range of the local ecotype (ecotype ranges determined using 95% confidence intervals) were assumed to be local recruits (red points). These samples were excluded from analyses.

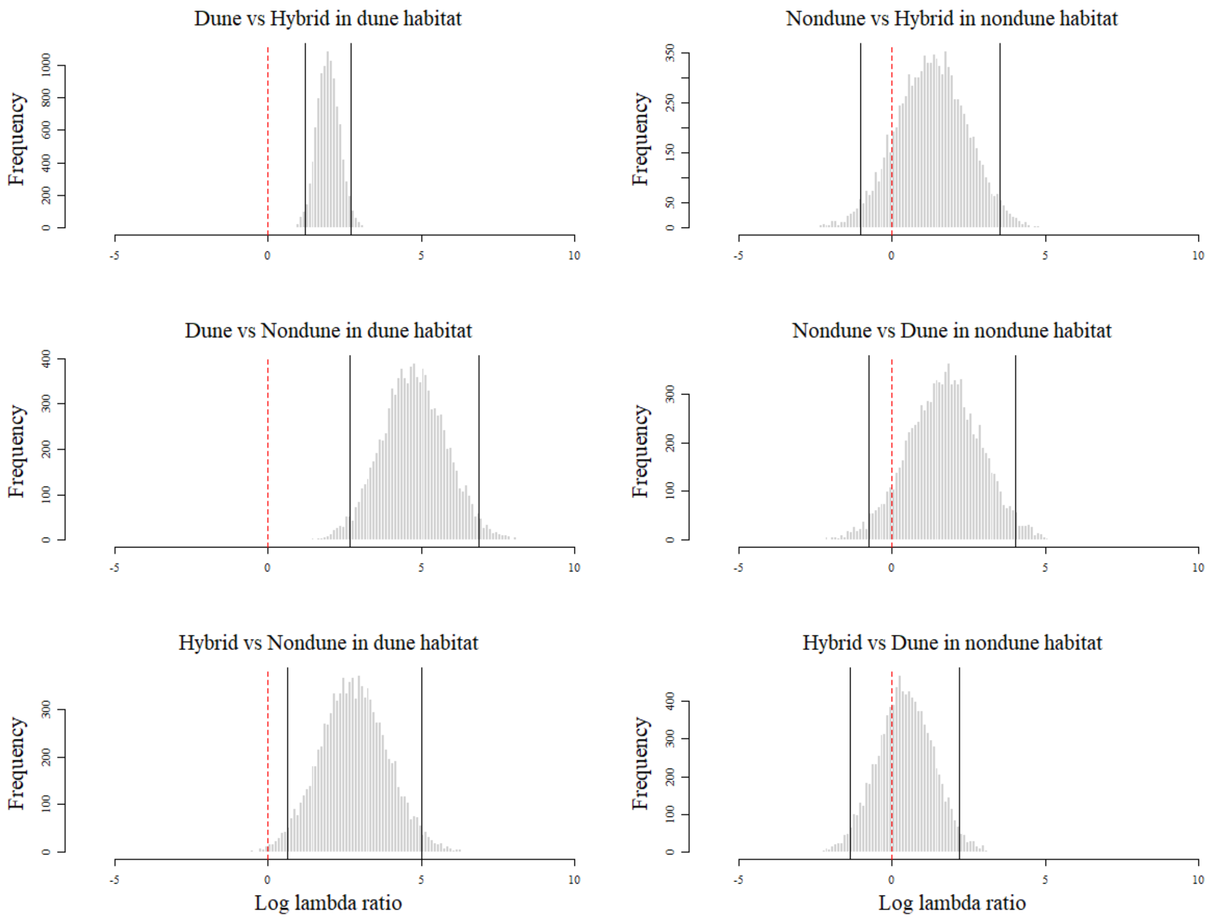

Figure S2: Parametric bootstrapping (10,000 replicates) used to calculate relative lambda (log ratio) between all source comparisons in both habitats. Vertical solid lines indicate 95% quantiles. Comparisons where zero (vertical dotted red line) is in the tail of the distribution indicate a significant difference between lambda estimates.

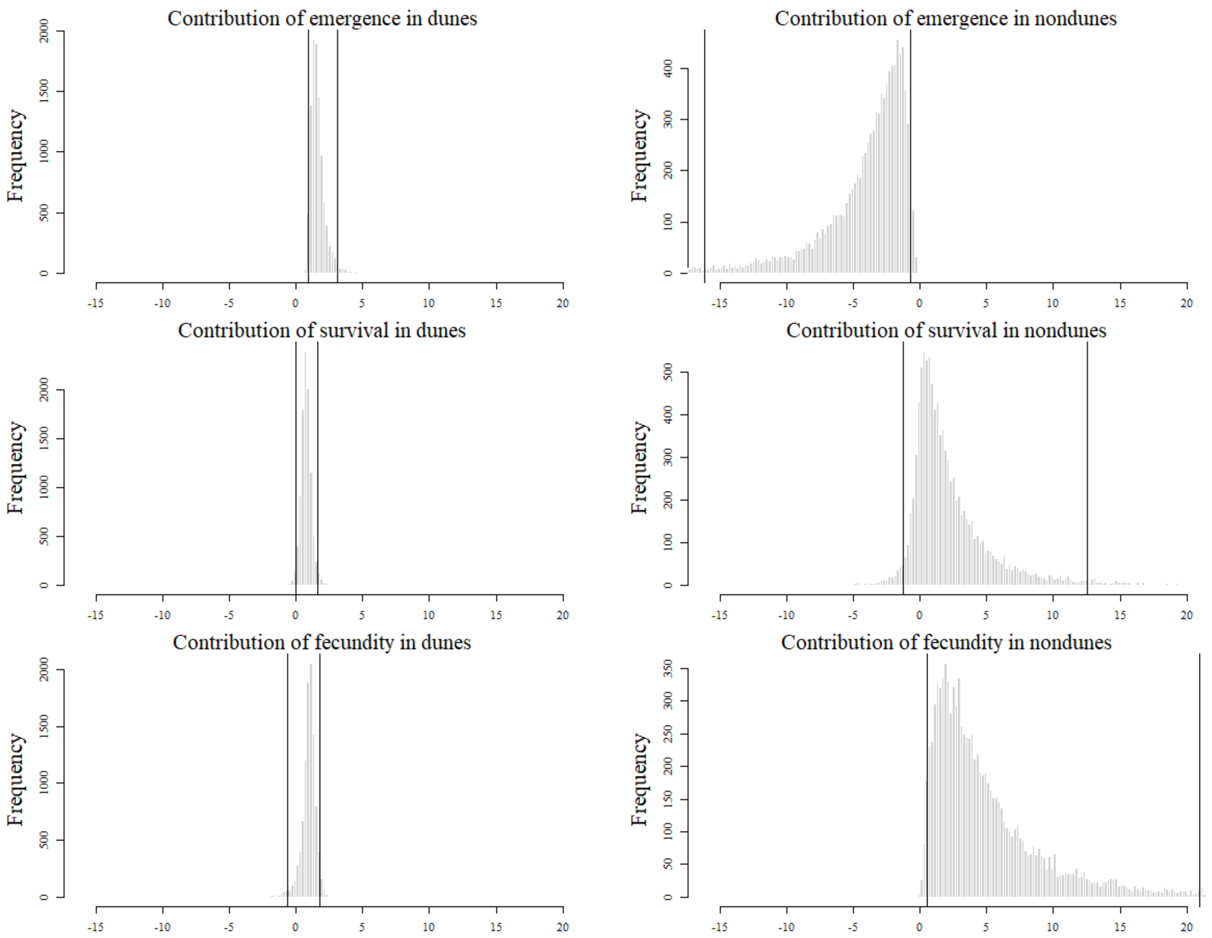

Figure S3: Distribution of fitness component contributions to differences in lambda between ecotypes (local – foreign) based on parametric bootstrapping (10,000 replicates). Contribution values are quantified as the change in a given fitness component value (local – foreign) multiplied by the sensitivity of lambda to that given fitness component. Vertical lines indicate 95% quantiles and were used to generate error bars in main figure 2.

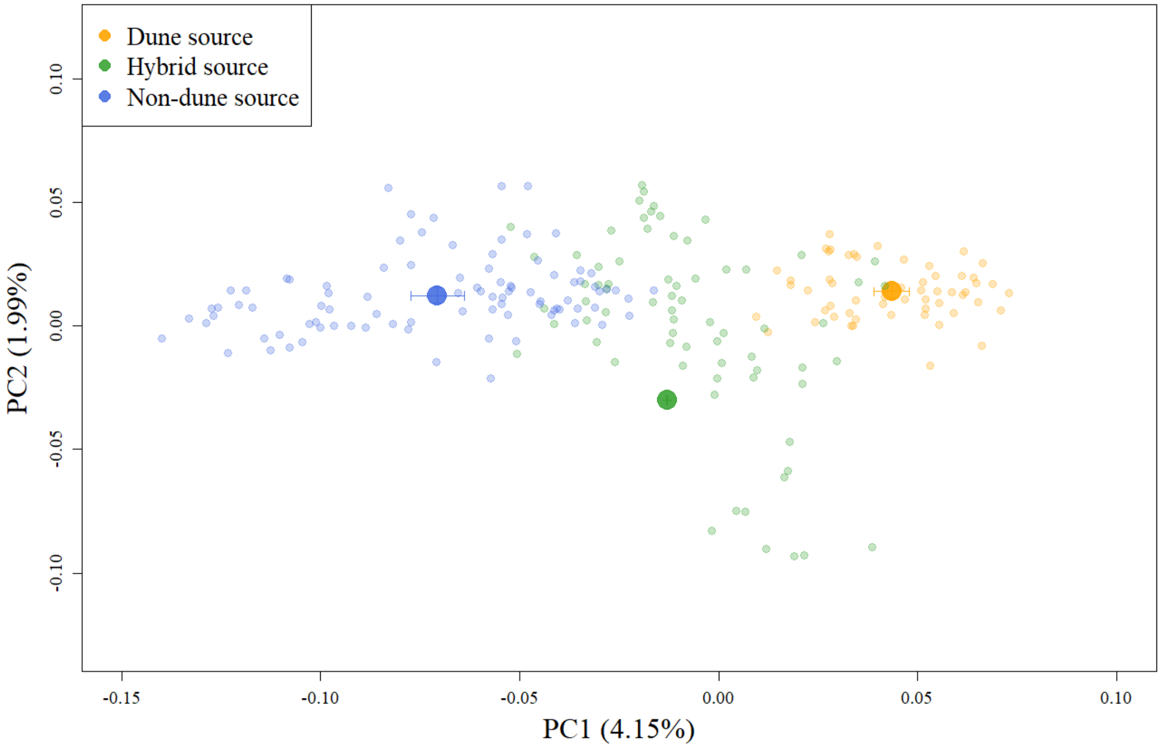

Figure S4: Principal component (PC) analysis based on 228 pre-selection samples genotyped at 12,214 SNPs. Scores for PC1 and PC2 for each sample (small points) as well as means (large points) and 95% confidence intervals for the average PC scores for each source are plotted. Note that mean hybrid score and associated confidence intervals are based on random sampling of individuals drawn in proportion to the number of each hybrid type planted in the field (see Supplemental Methods). This plot shows the genetic structure of pre-selection samples and the separation of sources along PC1.

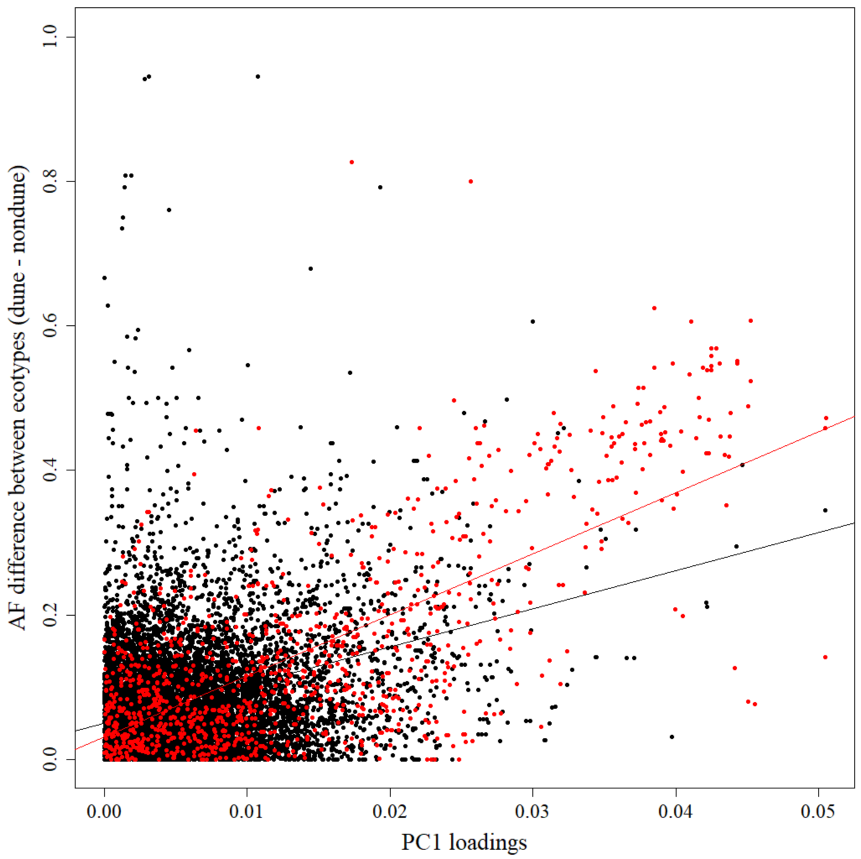

Figure S5: Absolute value of allele frequency (AF) difference between dune and non-dune ecotypes (post-selection samples from non-experimental plants; dune minus non-dune) plotted against the absolute value of PC1 loadings for all 12,214 SNPs. This plot supports ($R^2 = 0.155$, $p < 2.2e-16$; black line) the use of PC1 as an axis for separating samples based on genetic differences between ecotypes. The relationship between AF difference and PC1 loadings is driven more strongly by loci in inversions ($R^2 = 0.492$, $p < 2.2e-16$; red) compared to non-inversion loci ($R^2 = 0.040$, $p < 2.2e-16$; line not shown).

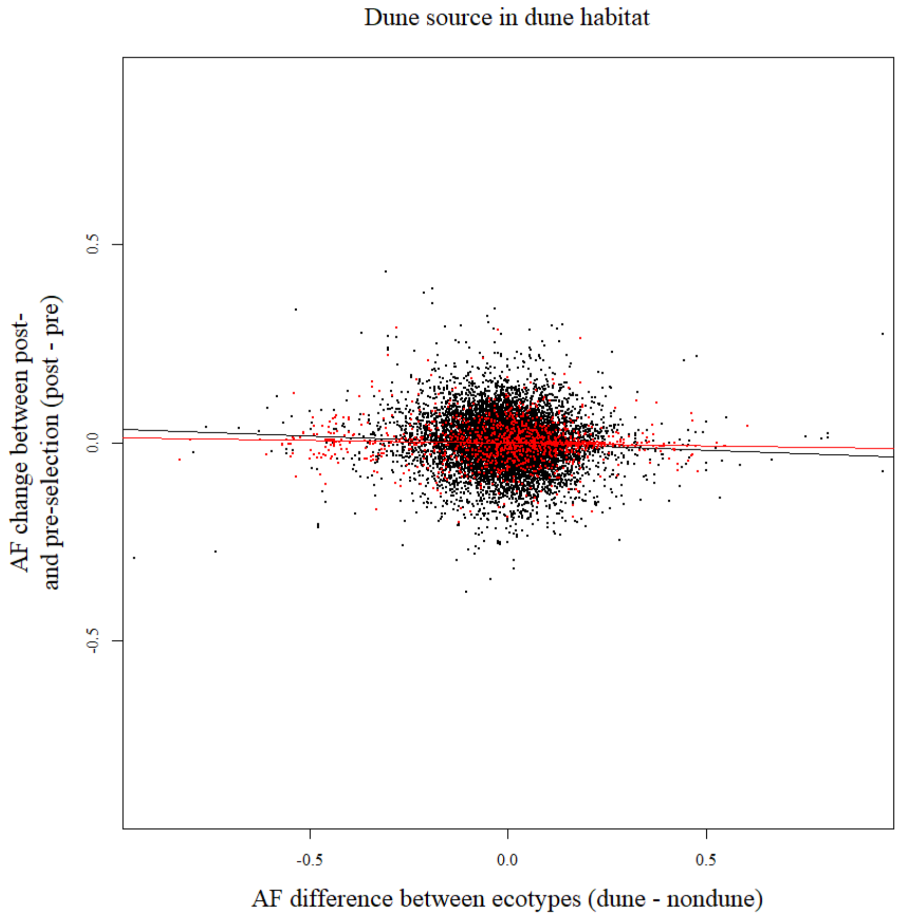

Figure S6: Relationship of allele frequency (AF) differences between ecotype (post-selection samples from non-experimental plants; dune minus non-dune) and allele frequency change during early life history (post- minus pre-selection) in the dune habitat in samples of dune source (r = -0.069, p = 0.064; all points). There was no significant relationship when considering inversion loci only (r = -0.052, p = 0.32; red), and a marginally significant relationship for non-inversion loci only (r = -0.074, p = 0.041; black points). P-values are based on one-sided tests using distributions from randomization tests (fig S7 B). Lines represent linear model fits (black = all points, red = inversion loci only).

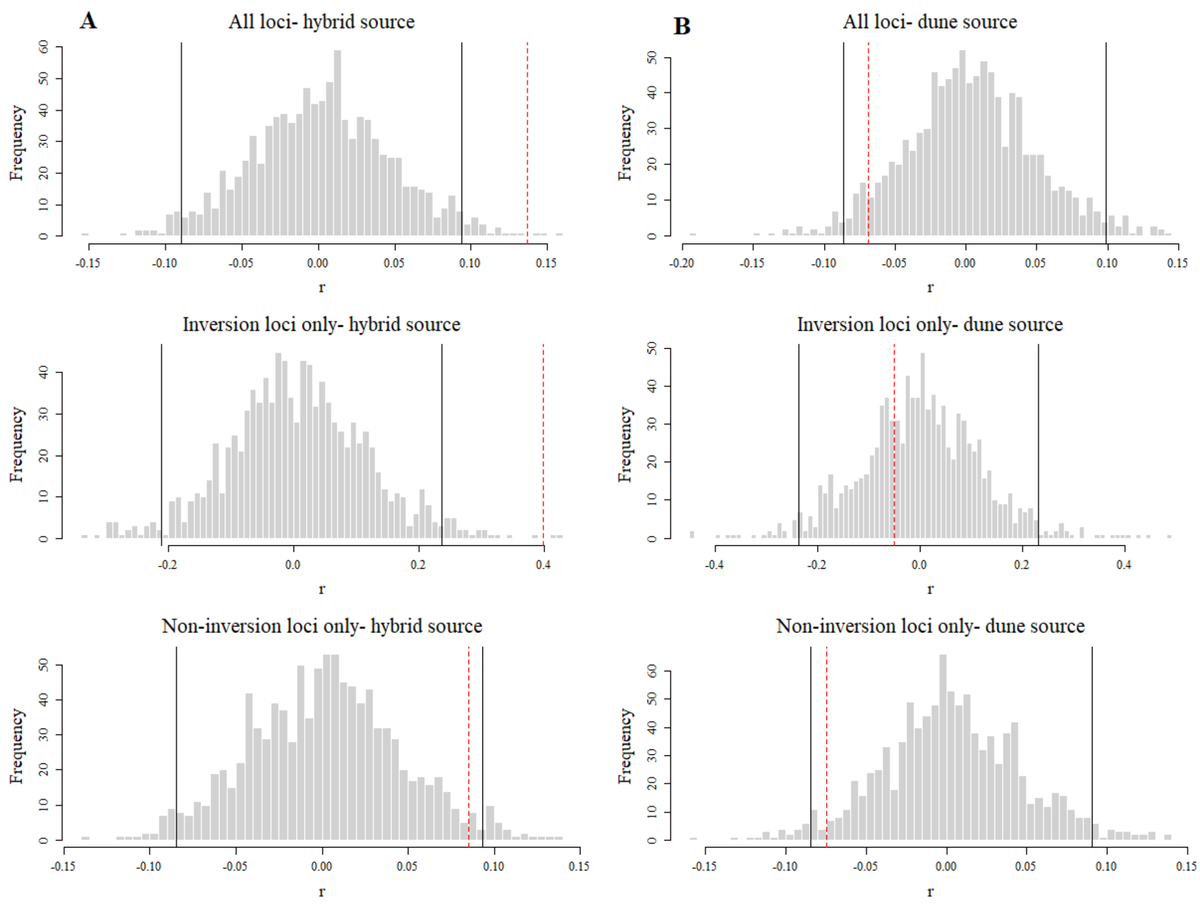

Fig S7: Randomization test to assess the significance of the patterns seen in A) fig 4 (hybrid samples) and B) fig S6 (dune samples). All samples used in the generation of A) fig 4 (non-experimental dune, non-experimental non-dune, pre-selection hybrid, post-selection hybrid) or B) fig S6 (non-experimental dune, non-experimental non-dune, pre-selection dune, post-selection dune) were randomly assigned a label corresponding to one of these 4 groups. Allele frequencies were calculated using the randomly assigned sample labels and differences were determined: dune minus non-dune and post-selection minus pre-selection. Pearson correlation coefficients (r) between these two differences were then calculated. This was done 1000 times and distributions of r are plotted. Black lines are 95% quantiles. Dotted red lines are r values for correlations using the true sample assignment; associated p-values were calculated using a one-sided test. The resulting p-values were qualitatively similar to parametric p-values from linear models.

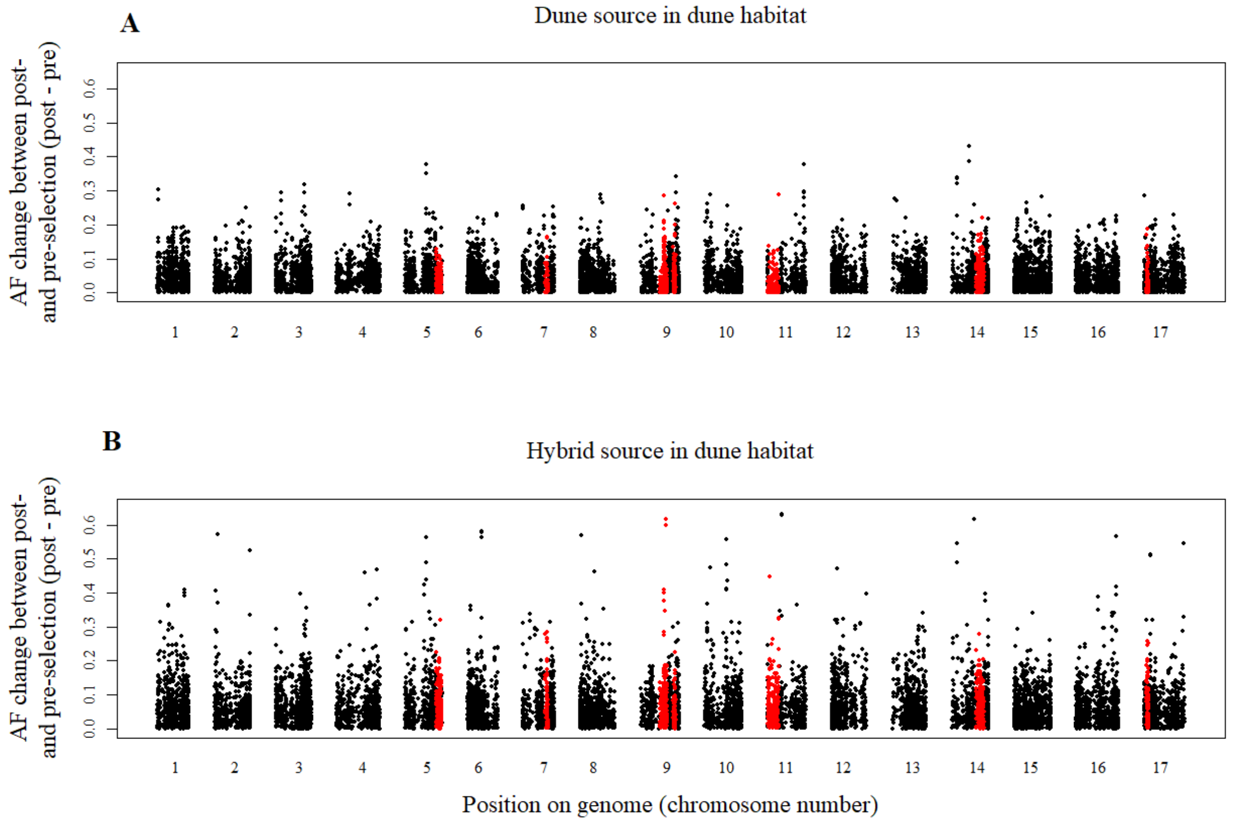

Figure S8: Genomic position of allele frequency (AF) change during early life history (post-minus pre-selection; absolute value plotted) in the dune habitat in samples of A) dune and B) hybrid source. Points are clustered by chromosome number. Red points indicate loci located in inversion regions. Note that non-dune source was excluded from this analysis due to low sample size.

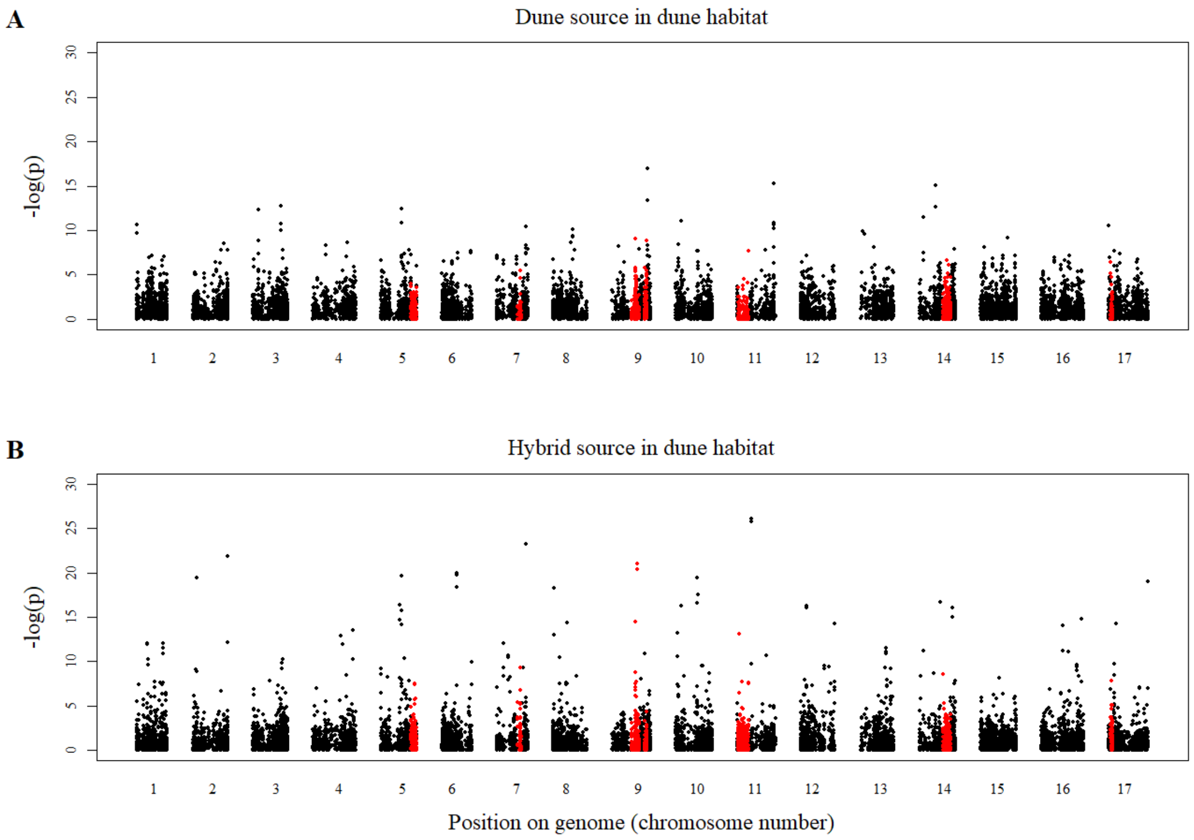

Fig S9: Genomic position of p-values (negative log) from models of allele proportions as a function of selection in the dune habitat in samples of A) dune and B) hybrid source. Points are clustered by chromosome number. Red points indicate loci located in inversion regions. Note that non-dune source was excluded from this analysis due to low sample size.

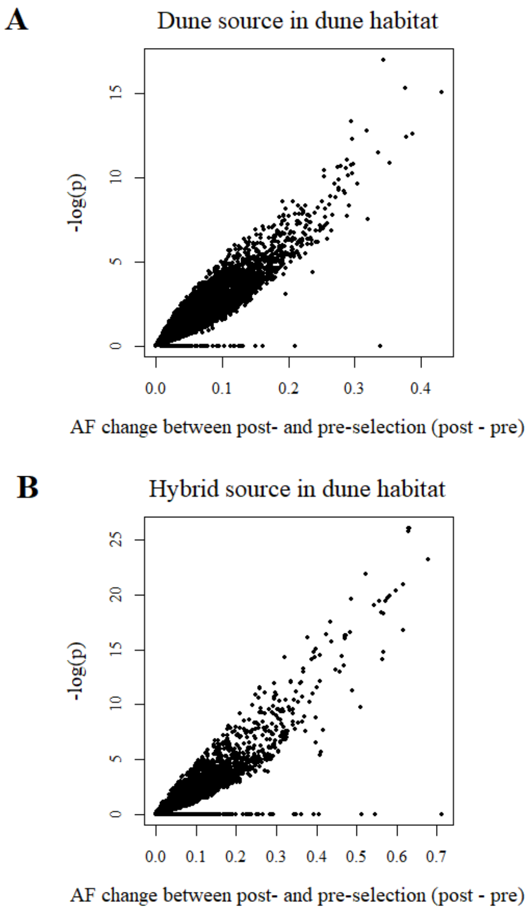

Fig S10: Relationship between allele frequency (AF) change during early life history (post- minus pre-selection; absolute value plotted) and p-values (negative log) from models of allele proportions as a function of selection in the dune habitat in samples of A) dune and B) hybrid source. Results from these two methods of assessing AF change are highly correlated: A) $R^2$= 0.84, p < 2.2e-16; B) $R^2$=0.76, p < 2.2e-16.

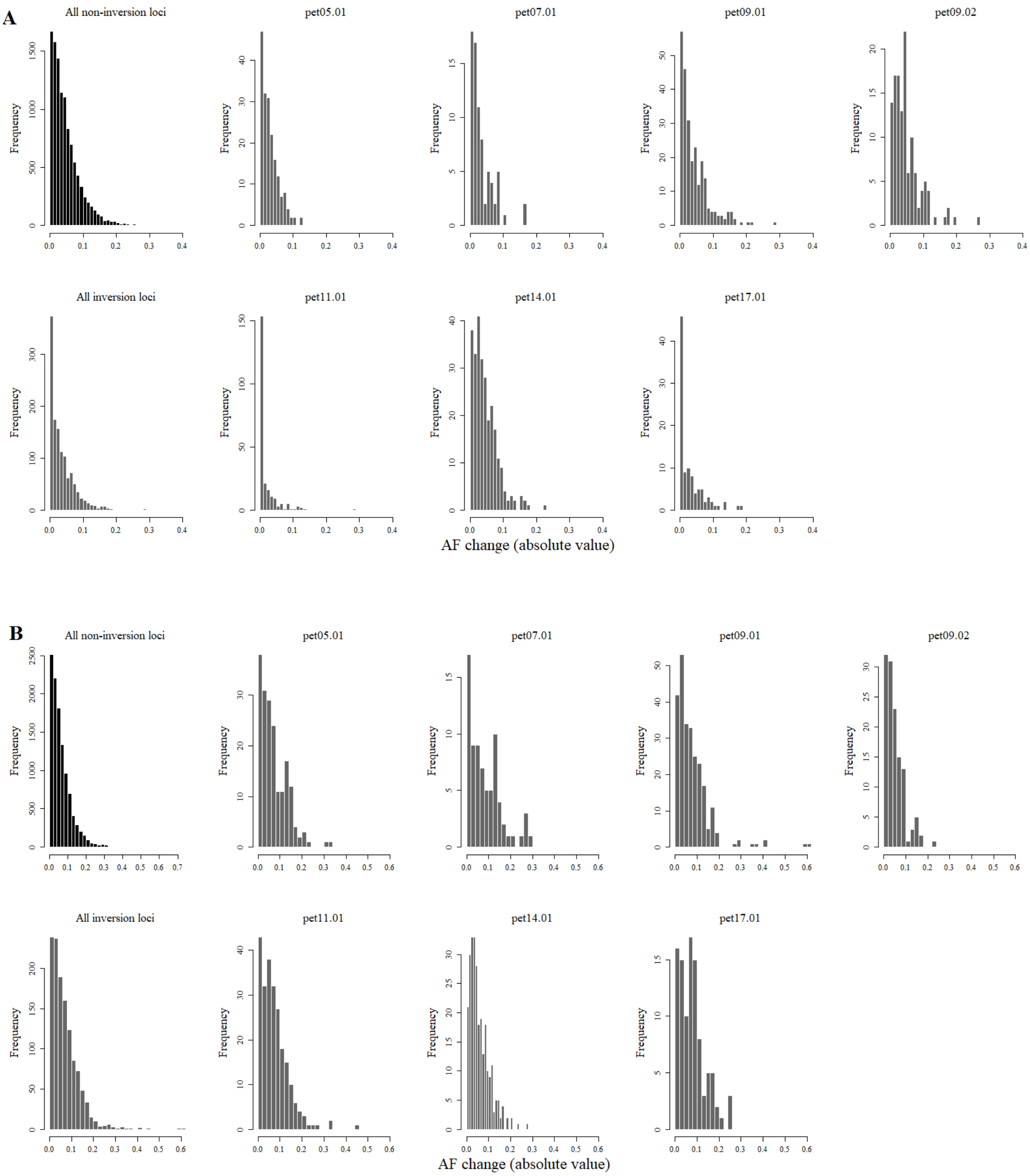

Figure S11: Distribution of allele frequency (AF) change during early life history (post- minus pre-selection; absolute value plotted) in the dune habitat for all non-inversion loci (black), all inversion loci (grey), and loci located in each individual inversion (grey) in samples of A) dune and B) hybrid source.
